# The hippocampal formation as a hierarchical generative model supporting generative replay and continual learning

**DOI:** 10.1101/2020.01.16.908889

**Authors:** Ivilin Stoianov, Domenico Maisto, Giovanni Pezzulo

## Abstract

We advance a novel computational theory of the hippocampal formation as a hierarchical generative model that organizes sequential experiences, such as rodent trajectories during spatial navigation, into coherent spatiotemporal contexts. We propose that the hippocampal generative model is endowed with inductive biases to identify individual items of experience (first hierarchical layer), organize them into sequences (second layer) and cluster them into maps (third layer). This theory entails a novel characterization of hippocampal reactivations as *generative replay*: the offline resampling of fictive sequences from the generative model, which supports the continual learning of multiple sequential experiences. We show that the model learns and efficiently retains multiple spatial navigation trajectories, by organizing them into spatial maps. Furthermore, the model reproduces flexible and prospective aspects of hippocampal dynamics that are challenging to explain within existing frameworks. This theory reconciles multiple roles of the hippocampal formation in map-based navigation, episodic memory and imagination.

## Introduction

During our lives, we continuously learn new skills and accumulate novel memories. A fundamental problem in both neuroscience and machine learning is preventing this novel acquired knowledge from interfering with past memories and causing *catastrophic forgetting* (McCloskey and Cohen, 1989). The hippocampus has long been considered important in solving this biological problem. According to the influential *Complementary Learning Systems* theory, it works as an episodic memory buffer that stores novel experiences rapidly and successively reactivates them offline (e.g., during sleep), in order to train a separate, semantic memory system located in the cortex, which gradually integrates and consolidates memories, preventing catastrophic forgetting (McClelland et al., 1995; O’Reilly et al., 2014).

The idea that the hippocampus stores and reactivates episodic memories offline has received empirical support from rodent studies during spatial navigation. In phases of wakeful rest during spatial navigation as well as subsequent sleep, the rodent hippocampus spontaneously generates (time-compressed) sequences of neural activations resembling those observed during behaviour, such as sequences of place cells corresponding to spatial trajectories experienced during navigation (Buzsáki, 2015; Foster, 2017), sometimes in reverse order (Foster and Wilson, 2006). These *internally generated hippocampal sequences* have biological significance as their disruption has been shown to impair memory-based navigation (Jadhav et al., 2012). These sequences have been called “replays” to highlight the fact that they might be interpreted as reactivations of previous experiences stored in an episodic memory buffer located in the hippocampus.

The idea of using an episodic memory buffer to store and replay experiences during learning is widespread in machine learning, as well. This “experience replay” method is especially effective for *continual learning*: when an artificial system has to learn multiple tasks sequentially. In most machine learning methods (e.g. those using neural networks), learning a novel task (task C) usually implies forgetting previously learned tasks (tasks A and B), because the memories of C interfere with those of A and B. Extending the neural net with experience replay mitigates this problem. Essentially, when the extended neural net learns tasks A and B, it stores a history of past episodes in a memory buffer. Then, while learning a novel task C, past episodes experienced during tasks A and B are intermittently sampled from the replay buffer and replayed thereby mitigating the risk of catastrophic forgetting. The functioning of this extended model has been often associated to the functioning of hippocampal “replay” during sleep (Kumaran et al., 2016; Mnih et al., 2015).

However, experience replay as commonly implemented in machine learning requires storing an unbounded quantity of verbatim memories and it is not clear whether and how the brain might implement something similar. An influential view describes the recurrent connections of the hippocampal area CA3 as an auto-associative memory network that stores static memory patterns (Treves and Rolls, 1992) or sequences of experiences (Cheng, 2013; Levy, 1996; Lisman, 1999). These networks could be set to spontaneously replay noisy versions of the stored memories, hence implementing something close to experience replay; but still they would require unbounded memory resources to store novel experiences during lifelong learning.

Furthermore, and importantly, increasing evidence challenges the view that the hippocampus simply stores and “replays” old memories (or noisy versions of old memories). In rodents, internally generated hippocampal sequences during sleep or wakeful rest can depict paths to future goal locations rather than only past trajectories (Pfeiffer and Foster, 2013), including optimized paths that had never been exploited by the animals (Igata et al., 2021); random trajectories, resembling Brownian diffusion (Stella et al., 2019); trajectories that have never been directly explored but reflect prior spatial knowledge (Dragoi and Tonegawa, 2013; K. Liu et al., 2018), such as shortcuts (Gupta et al., 2010) and future event locations (Ólafsdóttir et al., 2015); and *preplay* sequences of cells before a navigation episode, which are subsequently expressed during navigation (Dragoi and Tonegawa, 2011). In humans, internally generated sequences can re-order (non-spatial) item representations to reflect learned rules, rather than replaying the order in which they were actually experienced (Y. Liu et al., 2019).

In sum, internally generated hippocampal sequences during sleep or wakeful rest extend well beyond previous experiences and demonstrate prospective aspects that are difficult to reconcile with simple replay from a memory buffer (Foster, 2017; Penagos et al., 2017; Giovanni Pezzulo et al., 2017; Pezzulo et al., 2019). However, a robust theoretical framework unifying these findings has yet to emerge. Here we advance a novel theoretical proposal to explain flexible and prospective aspects of internally generated hippocampal sequences, within a statistical learning framework. We propose that the hippocampal formation functions as a *hierarchical generative model* that organizes sequential experiences (e.g., spatial navigation trajectories) into a set of mutually exclusive spatiotemporal contexts (e.g., spatial maps). Furthermore, we propose that internally generated hippocampal sequences stem from *generative replay*, or the resampling of fictive experiences from the generative model, rather than from a verbatim replay of memories from a buffer.

This new theory is based on four main assumptions. First, that the hippocampal formation encodes experiences by *learning a hierarchical generative model* of observed data. The proposed hierarchical model has three layers, see Figure 1A. The first hierarchical layer forms latent codes for individual *items* of experience (e.g., the animal’s spatial location) inferred from sensory observations. These item codes are initially deprived of context. However, the second and third layers organize items into coherent spatiotemporal contexts. Specifically, the second layer forms latent codes for *sequences* within which the lower-level items are embedded (e.g., the animal’s spatial trajectory), supporting their sequential processing – hence establishing the hippocampus a “sequence generator” (Buzsáki and Tingley, 2018). The third layer forms latent codes for structured *maps* that cluster experiences into mutually exclusive spatial or relational contexts (e.g., the maze the animal is currently in versus alternative mazes). The hierarchical structure is key for the functioning of the hippocampal model, as it provides guiding principles (or “inductive biases”) to organize individual *items* of experience in meaningful ways: namely, as *sequences* (temporally ordered items) and then *maps* (spatially clustered sequences).

**Figure 1.**
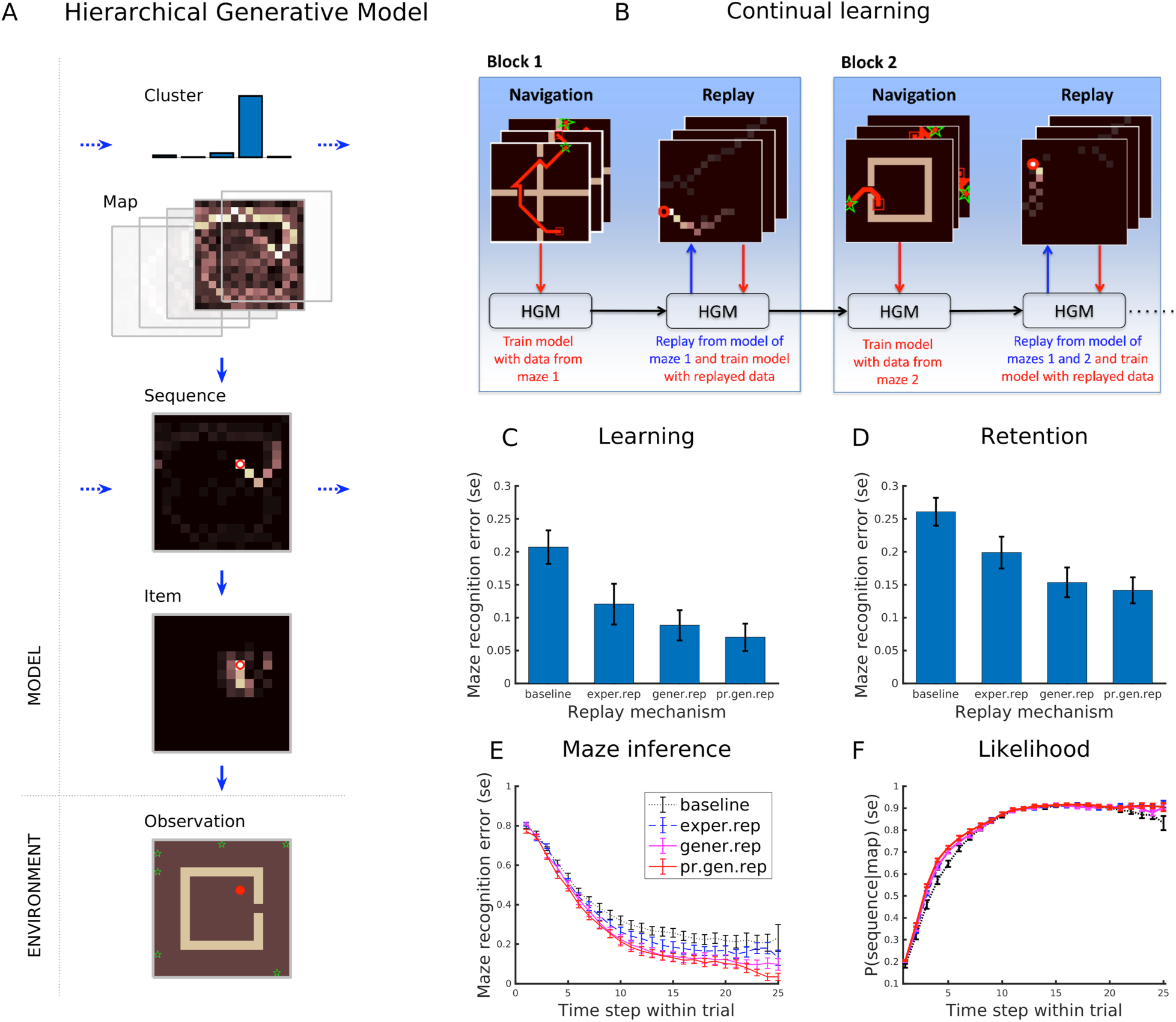
Hippocampal generative model. (A) Components of the hierarchical generative model, from top to bottom: clusters / maps, sequences, items and observations. The red circle denotes the agent’s current position. Colour codes correspond to probabilities, from high (white) to low (black). See the main text for explanation. The current implementation focuses on spatial navigation, where items, sequences, and maps may correspond to place cells, place cell sequences and 2D spatial maps, respectively. However, the same model could also apply to other domains, where the structure of maps could be different and even include factorizations (e.g., separate maps for objects or locations). (B) Structure of the continual learning experiment. We compare four learning agents having the same hierarchical generative model but different replay mechanisms: no replay, experience replay, generative replay and generative prioritized replay. The agents’ task consists in learning five mazes, each presented in a different block. For each block, the learning agents receive 20 trajectory data from one single maze, and use them to train their hierarchical generative model. Between blocks, the learning agents can replay experiences in different ways; see the main text. (C-F) experimental results (means and standard errors of n=16 replicas of each agent). Baseline: baseline agent. Exper.rep: experience replay agent. Gener.rep: generative replay agent. Pr.gen.rep: prioritized generative replay agent. (C) Performance of the four agents during learning (i.e., inference of the correct maze). In this and all the next plots, error bars denote standard error. Please note that chance level is 20%. (D) Same as C, but measured during a separate (retention) test session executed after the agents experienced all five blocks. (E,F): Dynamics of inference. (E) Same as D, but as a function of time step within trials (for 25 time steps). (F) Accuracy of the reconstruction of sequences, as indexed by the likelihood of the sequence given the correct map. See the main text for explanation.

The second assumption is that the hierarchical generative model serves a fundamental function: binding experiences into coherent spatiotemporal contexts (Buzsáki and Moser, 2013; Yonelinas et al., 2019). This notion stems directly from the fact that in the hierarchical model, sets of item and sequence codes at the lower level (which correspond to individual experiences) are organized into separate maps (which correspond to mutually exclusive relational contexts). Hence, experiences that share surface similarities, such as running in two similar corridors, can engage different map codes in the generative model - and sometimes, different sets of neurons. The possibility to bind sets of individual experiences to separate relational contexts is a key feature of episodic and autobiographical memory (Tulving, 2002). From a computational perspective, the possibility to group experiences into separate contexts might be key to preventing catastrophic forgetting of episodes. This idea is in keeping with a large body of literature showing that context learning improves robustness and generalization (Collins and Frank, 2013; Gershman et al., 2010; Heald et al., 2021; Sanders et al., 2020; Stoianov et al., 2015) and that architectures that use different, non-overlapping units for different tasks, such as cloned HMM (Rikhye et al., 2020) and others (Masse et al., 2018) are especially effective for continual learning.

The third assumption is that internally generated hippocampal sequences are manifestations of the inferential dynamics of a hierarchical generative model rather than “replays” of old experiences from a memory buffer or an associative memory (Dudai, 2012; Foster, 2017; Ólafsdóttir et al., 2018; Pezzulo et al., 2014). Therefore, internally generated hippocampal sequences can support both the re-enactment of experienced events (and their context) and the imagination of novel, never experienced events having a similar statistical structure as the events learned by the generative model.

Finally, and importantly, we assume that a key function of internally generated hippocampal sequences is learning and optimizing the hippocampal generative model for future use. Specifically, we posit that the hippocampal generative model supports *generative replay*: a method for continual learning that uses fictive experiences resampled from a generative model to self-train the model that generates them (as well as to train separate models or controllers). Previous research has established that generative replay is appealing from a computational perspective, since it can prevent catastrophic forgetting and (unlike experience replay) does not require unbounded memory resources to store previous experiences (Mocanu et al., 2016; Shin et al., 2017a; van de Ven et al., 2020). Here, instead, we use the concept of generative replay to explain flexible and prospective aspects of internally generated hippocampal sequences, which are difficult to reconcile with previous theories.

The remainder of this article comprises two main sections. In the first Section, we illustrate the functionality and computational efficacy of the novel hypothesis, using a continuous learning task that requires learning multiple mazes. We compare the learning performance of different artificial agents, with or without replay mechanisms or a deep hierarchical structure. To preview our results, we show that the combination of generative replay and hierarchical structure affords the most effective continual learning. Agents using generative replay learn better spatial models compared to those without replay and (in the most challenging cases) to agents with perfect experience replay, which is biologically infeasible. Furthermore, endowing agents with a deep hierarchical structure increases learning performance and permits encoding different mazes into separate spatial maps, hence providing resistance to catastrophic forgetting.

In the second Section, we will illustrate the biological relevance of the novel hypothesis and its capability to explain empirical findings. Our simulations will reproduce the fact that replay dynamics are sensitive to goal information (Ambrose et al., 2016), that their statistics reflect preconfigured neural patterns more than navigational experience (K. Liu et al., 2019) and that they follow Brownian diffusion-like random trajectories even if the animal’s trajectories during navigation were not Brownian (Stella et al., 2019).

## Results

### Continual learning of multiple spatial maps with generative replay

To assess the computational efficacy of generative replay, we designed a continual learning task consisting of learning multiple spatial mazes from (simulated) navigation trajectories. The continual learning task consists of five 15×15 mazes (shown in Figure 3A) experienced sequentially, in five different blocks. To render the learning experiment challenging, we selected five mazes that have large portions of spatial structure in common; the only difference between the mazes is that the walls are in different locations.

Our proposed approach solves the task by learning a hierarchical generative model of its spatiotemporal (navigation) experiences (Figure 1A-B). Our simulations will show that the learned model permits inference of the present maze and generation of fictive replay trajectories from the same maze, which in turn can be used to improve the generative model itself and to predict future navigational experiences.

We compare four agents that learn the hierarchical generative model using different replay methods: (1) a “baseline” agent with no replay; (2) an ideal “experience replay” agent that replays previous experiences from a memory buffer to train the generative model; (3) a “generative replay” agent that replays experiences from its learned generative model to train the same generative model; (4) a novel variant of generative replay, which we call “prioritized generative replay”, which is the same as the previous agent, but uses a different replay method, which prioritizes surprising experiences. Please see the Supplementary materials for the pseudocode of the four agents.

Comparing the three replay agents with the baseline agent is useful to illustrate the advantages of replay. Further, comparing the two generative replay agents with the ideal experience replay agent is useful to assess the advantages and disadvantages of replaying trajectories from a generative model that is necessarily imperfect and can produce out-of-sample data, compared to a (perfect) model of previous experiences. Note that we call the experience replay agent “ideal”, because storing a lifetime of verbatim memories is biologically infeasible.

Below we introduce the agents’ hierarchical generative model (which is the same for all agents) and their training procedures (which is different for each agent).

#### Hierarchical generative model

All agents are endowed with the same hierarchical generative model, shown in Figure 1A (see the Methods section for details). The generative model is composed of three layers of hidden states, encoding *items*, *sequences* and *maps*, respectively, as well as a bottom layer encoding the present *observation*. We describe the model from the bottom-up, beginning with observations.

At each moment, the agent receives an *observation* about its current spatial location in the maze (or an item in a spatial location). Note that the agent never observes full trajectories, but instead receives one data point at a time.

The first hidden layer of the hierarchical model encodes *item* codes, putatively corresponding to place cells. Note that while in Figure 1A *observations* and *items* are plotted similarly in grid coordinates, they are distinct elements: observations correspond to model inputs and are coded as one-hot vectors, whereas items are hidden (or latent) states of the generative model and correspond to probability distributions. The model does not receive item codes as inputs but rather infers them using a scheme analogous to predictive coding (Friston, 2005; Rao and Ballard, 1999), by integrating four sources of information: the current bottom-up observation, top-down predictions from the higher hierarchical layers, lateral information from the previous sequence of items, and a model of movement dynamics (see the Methods section).

The second hidden layer of the hierarchical model encodes *sequences* of items, which are inferred by considering three sources of information: lateral information from the sequence inferred at the previous time step, bottom-up information from the last inferred item, and top-down information from the currently inferred map.

The third hidden layer of the hierarchical model encodes *maps* (i.e., clusters of items) using a probabilistic mixture model: a way to represent the fact that different sequences of items can belong to different, mutually exclusive clusters (i.e., different spatial trajectories can belong to different spatial maps). Each map corresponds to a probability distribution over an arbitrary structure. Please note that this notion is generic: a map is simply a vector of items, each with an associate probability, with no fixed semantics. However, here we use the generic notion of map to denote a hippocampal spatial map, in which the probability distribution corresponds to an occupancy distribution over locations and the structure corresponds to adjacency relations between locations. Figures 1 and 3 provide several examples of spatial maps (plotted as 2D grids for ease of interpretation) that emerge in our model. During learning, the maps acquire information about the relations between its elements (items), based on two factors: *input statistics* and *structural priors*. The former (input statistics) correspond to the sequences of observations about the agent’s locations in the maze that the model receives during learning. The latter (structural priors) depend on the way the model infers the next item. During inference of the next item, the model makes two assumptions: first, that the locations of successive items change gradually (and hence there are only “small gaps” between consecutive locations) and seconds, that there is no preferred directionality between successive items; see equation 8 of the Methods section for a formal description of these assumptions. This implies that the model is preconfigured (i.e., has *inductive biases*) with spatial-topological constraints, such as the fact that sequences of items (trajectories) have few spatial discontinuities and can be bidirectional. Both input statistics and structural priors contribute to gradually turn naive vectors of items into maps that encodes a space’s topological information and adjacency relations between locations, as commonly hypothesized for hippocampal maps (Dabaghian et al., 2014).

Furthermore, maps do not simply represent items in spatial locations but also their *behavioural relevance*. This is because the items have associated probability distributions that are updated using visitation counts during both real and fictive experiences (explained in detail later) and come to reflect the future expected occupancy – or empirical priors in Bayesian parlance (but note that using other methods, such as algorithmic complexity (Donnarumma et al., 2016; Maisto et al., 2016, 2015) and successor representations (Gershman, 2018; Stachenfeld et al., 2017), would produce similar results as visitation counts). This expectation is typically zero at locations that cannot be crossed (e.g., walls) and high in frequently visited locations, such as goal sites and other behaviourally relevant locations, such as junction points or bottlenecks (Aoki et al., 2019). The fact that maps encode behavioural relevance reflects the idea that some aspects of value are incorporated into the hippocampal map and influence replay statistics (Bhattarai et al., 2020). This is an aspect that we will discuss below, in the section on “Biological interpretation of generative replay and supporting evidence”.

Finally, the third layer of the hierarchical model continuously infers which map the agent is currently in. For this, the model maintains a (categorical) probability distribution over maps (or clusters) and updates it whenever it receives new observations, real or fictive (i.e., produced by generative replay). The map having the highest probability is the currently inferred one; for example, in Figure 1A, the currently inferred map is the fourth. Crucially, at each moment in time, only the currently inferred map is updated during learning and used for generative replay. This selective learning strategy permits acquiring well-differentiated maps and is key to counteract catastrophic forgetting.

In this simulation, the generative model is provided with five clusters. This is done to facilitate the analyses of the correspondences between maps and mazes, under the hypothesis that each cluster will “specialize” in one of the five mazes. Later, we will generalize this approach to an arbitrary number of clusters, using a nonparametric method that automatically selects the best number of clusters, given the agent’s observations (see the section on “Nonparametric model”).

#### Training procedure

The four learning agents are trained in the same way *within* blocks: they receive 20 spatial trajectories from the maze they are currently in as input (experiencing each trajectory just once in pseudorandom order and one location at a time) and use these data to train their generative models.

The agents learn the five mazes in different blocks (mazes were selected randomly without repetitions for each agent and block). During a block, each of the four learning agents receives the same batch of 20 spatial trajectories from the current maze, and uses these data to update their own hierarchical generative model. The trajectory data are generated by a separate model-based Bayesian controller that, for each trial and maze, moves between a randomly selected origin and goal location, by following a near-optimal trajectory (Stoianov et al., 2018).

What distinguishes the four agents is how they update their generative model *between* blocks, see Figure 1B. The first (baseline) agent does not update the generative model between blocks. The other three agents use replay to further train their generative models offline: they replay 20 trajectories (to make them commensurate with real experience; see also the control simulations in the Methods section) and use these fictive data to update their generative models, in exactly the same way they do with real trajectories. However, each agent uses a different replay mechanism. The second agent uses *experience replay*: it replays trajectories randomly sampled from a full experience history. Note that this is an idealized agent that uses unbounded resources to store all its memories without loss; its performance should be seen as an upper bound to the capabilities of any biologically plausible agent that performs experience replay with bounded resources. The third agent uses *generative replay*: it resamples trajectories from its currently inferred map, i.e., the one corresponding to the cluster with maximum probability (note that as the probabilities are continuously updated, successive replays can be generated from different learned maps). The fourth agent uses *prioritized generative replay*: it samples trajectories from its generative model, but using *prioritized maps* to initialize a replay instead of the standard maps used by the third agent. Prioritized maps are auxiliary maps (one for each cluster) that encode the average *surprise* that the agent experienced at each location during navigation, rather than visitation counts. Average surprise accumulates (Shannon) surprise derived from the probabilities encoded in the maps, for each location. Using prioritized maps permits resampling trajectories corresponding to the portions of the maze that were sampled less frequently during navigation and hence might be coded imperfectly, similar to prioritized sweeping in reinforcement learning (Moore and Atkeson, 1993), hence potentially explaining why novel sequential activations were replayed more frequently after exposure (Dragoi and Tonegawa, 2013; K. Liu et al., 2018).

#### Simulation results

The results of the continual learning experiment are shown in Figure 1C-F. Figure 1C shows the agents’ learning error, measured as their ability to correctly infer what maze generated their observations (i.e., which maze they were in) – which in Bayesian terms, is a form of *latent state (maze*) inference (Fuhs and Touretzky, 2007; Penny et al., 2013; Sanders et al., 2020). To compute learning error for a trial we first establish the agent’s best matching cluster for each maze at the end of learning, which we consider to be a “ground truth,” given it is the cluster the agent would select in the limit with infinite data. Note that we select the single highest probability cluster despite the fact that in some cases, and most notably when using the non-parametric model, it is possible that several clusters have non-negligible probabilities for a given maze. Next, we count for each step how many times the agent correctly inferred the “ground truth” cluster, after the first 10 observations of each trial (but we exclude the first 5 trials of each block from this analysis). All the agents showed a good degree of contextual learning and a good memory of the old mazes after learning novel mazes. This capability to counteract catastrophic forgetting stems directly from the possibility to organize experiences into mutually exclusive contexts or maps and is therefore common to all the agents. However, the two agents using generative replay and prioritized generative replay outperformed the baseline agent (t=3.47, s.d.=0.10, df=30, p=0.002 and t=4.18, s.d.=0.09, df=30, p<0.001, respectively) and showed no difference from the ideal experience replay agent with perfect memory (t=0.83, n.s. and t=1.35, n.s., respectively). Finally, we found no significant differences between generative replay and prioritized generative replay agents (t=0.587, n.s.).

Figure 1D shows the same learning error, but computed during a separate (retention) test session executed after the agents experienced all five blocks. During the retention test session, we reset the cluster probabilities after each trial; hence the agent treats each new trajectory as independent. The retention test is therefore more challenging than the learning task, because the agents cannot accumulate trial-by-trial evidence about the maze they are currently in. Furthermore, learning novel mazes could lead to catastrophic forgetting of previous mazes. Again, the two agents using generative replay and prioritized generative replay outperformed the baseline agent (t=3.47, s.d.=0.09, df=30, p=0.002 and t=4.13, s.d.=0.08, df=30, p<0.001, respectively) showed no difference from the ideal experience replay agent (t=1.37, n.s. and t=1.84, n.s., respectively). Finally, we found no significant differences between generative replay and prioritized generative replay agents (t= 0.401, n.s.).

Figures 1E and 1F characterize the dynamic effect of evidence accumulation *within* trials. They plot the performance of the agents during the test session during the first 25 time steps of each trial, during which agents gradually accumulate information about both the maze and the trajectories they observe (recall that agents receive one observation at a time and never observe entire trajectories). Our results show that as they accumulate more evidence within a trial, all agents demonstrate increasing maze inference accuracy (Figure 1E) as well as increasing likelihood of generated sequences (Figure 1F). The latter is equivalent to reconstruction accuracy, which is a performance measure widely used to train autoencoders and generative models in machine learning. To compare the maze inference accuracy of the agents over time, we aggregated the results shown in Figure 1E into five intervals of five time steps each (1-5, 6-10, etc.). We found that the two generative replay agents (averaged) outperform the baseline agent across all five intervals (interval 1: t=2.658, sd=0.032, df=30, p=0.012; interval 2: t=2.688, sd=0.082, df=30, p=0.012; interval 3: t=3.931, sd=0.083, df=30, p=0.000; interval 4: t=3.606, sd=0.086, df=30, p=0.001, interval 5: t=3.080, sd=0.125, df=30; p=0.004) and also the experience replay agent, but only in the fifth interval (t=2.605, sd=0.087, df=30; p=0.014).

The benefits of generative replay and prioritized generative replay can be further appreciated by analysing the clusters they form during learning (see Figure 2). As shown in Figure 2A, all the agents tend to select one cluster for each maze, but the two agents using generative replay and (especially) prioritized generative replay do this more systematically. The analysis of the cluster-maze “confusion matrices” of the four agents (Figure 2B) shows that the two agents using generative replay and prioritized generative replay disambiguate more clearly between the five mazes they experienced.

**Figure 2.**
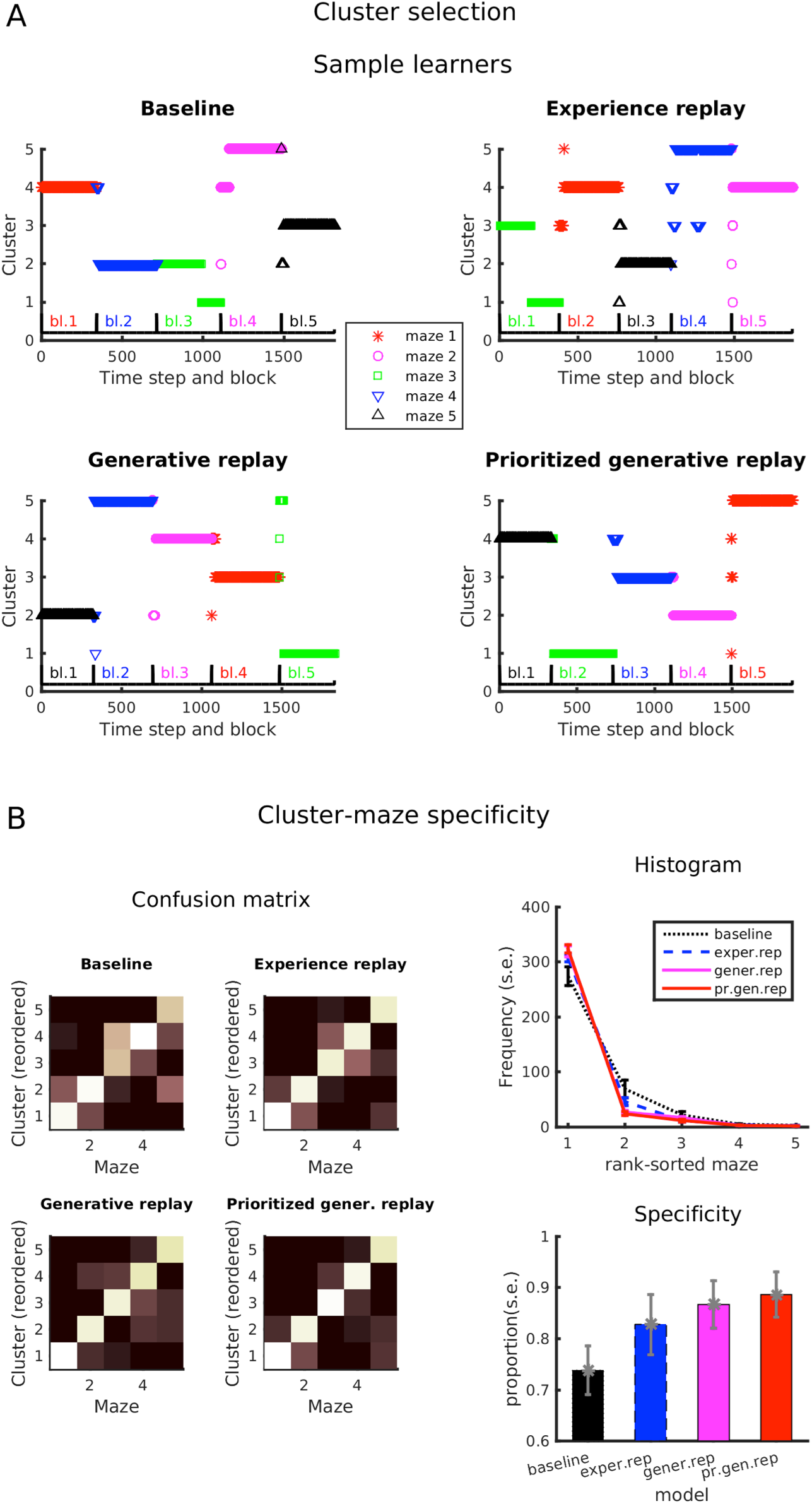
Cluster specificity during the continual learning experiment. (A) Cluster selection during navigation in sample replicas of each agent. The abscissa is double-labelled with time step (below) and colour coded blocks (above). The ordinate indexes the clusters (1 to 5) selected during the blocks. The five mazes correspond to different colours and symbols. In most cases, agents select different clusters for each block (especially after offline replay). (B) Analysis of cluster-maze specificity across all n=16 replicas of the four learning agents. Left: average cluster-maze confusion matrices. Right: Average frequency of cluster selection across clusters as a function of cluster-specific maze-preference rank (top) and proportion of the top-rank cluster selected during navigation (bottom).

**Figure 3.**
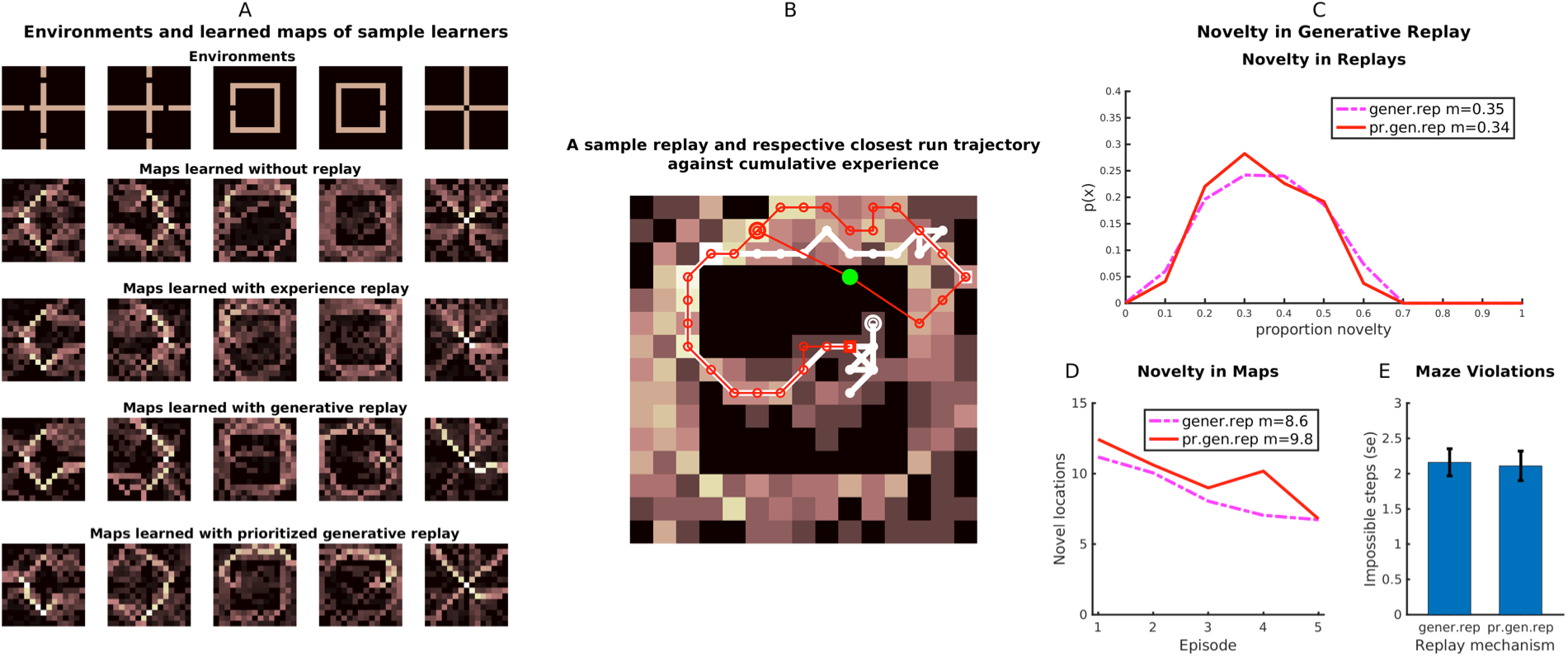
Examples of maps and replays of generative replay agents. (A) The five mazes used in the simulations (top) and some example maps learned by the four agents considered in this study. Colours represent probabilities, from white (high) to black (low). (B) Example novel trajectory produced during generative replay (red) superimposed on the closest trajectory (white) experienced during navigation in the third map shown in panel A. The novel trajectory starts from the double circle on the top-left and passes through several adjacent points, while sometimes making longer “jumps” in the map. Please note that this is an “impossible” novel trajectory that crosses a wall (in correspondence to the green point). (C) Quantification of the proportion of novel map locations visited during generative replay. This panel shows the extent to which the trajectories replayed by the generative replay and the prioritized generative replay agents differ from the most similar trajectories experienced during navigation (average = 35% and 34%, respectively). (D) The frequency of novel locations produced during generative and prioritized generative replay events (average = 8.6 and 8.9, respectively). (E) Average number of impossible locations visited (violations of maze constraints) during generative replay.

To assess the generality of these findings, we ran various control simulations, in which we tested each agent in the same Learning and Retention tests, while varying three parameters: the number of replays, the displacement coefficient parameter (*c*) that controls sequence continuity (Supplementary Table S1), and the number of training trials, i.e., the number of spatial trajectories that the agents experience during spatial navigation (Supplementary Figure S6). All our simulations show a consistent advantage of the three replay agents over the no-replay baseline, which is especially prominent when the replay agents experience a small number of trajectories as inputs (1 to 16) but can replay multiple times (20 times), see Supplementary Figure S6A-B. Interestingly, the two generative replay agents consistently outperform the experience replay agent when there are few training trials (1 to 16), whereas the differences between the four agents diminish as the number of training trials increases, see Supplementary Figure S6.

In sum, the comparison between the three replay agents and the baseline agent shows that replay improves the continual learning of multiple mazes. The comparison between the two generative replay agents and the ideal experience replay agent shows that despite the generative replay agents’ imperfect models, they perform at the same level of, and in some tests even outperform, an experience replay agent with perfect memory.

A possible reason for the effectiveness of generative replay is that it uses learned maps of the mazes (Figure 3A) to generate novel - but largely plausible - trajectories during replay (Figure 3B). The maps learned by the models, shown in Figure 3A, encode spatial-topological information consistent with the actual mazes and the *probability* of items at given locations; probabilities are colour coded, with white (black) corresponding to higher (lower) probabilities. These probabilities are important as the replay events follow high probability paths. Figure 3B shows an example novel trajectory produced by generative replay (in red), superimposed over an actually experienced trajectory (in white). On average, the trajectories produced during generative and prioritized generative replay differ from the most similar trajectories experienced during navigation by 35% and 34%, respectively (Figure 3C) and they include 8.6 and 8.9 novel locations that the agent never experienced during navigation (Figure 3D).

From a machine learning perspective, the possibility to generate novel trajectories might seem like a limitation rather than a feature, given that such trajectories are sampled from an imperfect generative model and are therefore less faithful to underlying world dynamics compared to experience replay. Furthermore, generative replay might sometimes generate “impossible” trajectories that violate the physical structure of the mazes (e.g., pass through walls), as shown in Figure 3B. A recent experiment reports that during hippocampal replay, impossible trajectories are present but much rarer compared to novel meaningful sequences (Widloski and Foster, 2022). Similarly, our results show that generative replay produces only a minority of impossible events (Figure 3E). This suggests that generating novel trajectories during replay is advantageous for learning, rather than hindering it.

One reason why generative replay produces plausible novel events is that the model’s inferential process is preconfigured with spatial-topological constraints (or *structural priors*), such as the fact that trajectories have only small gaps and can be bidirectional. By generating novel trajectories that reflect such spatial-topological constraints, generative replay and prioritized generative replay agents can effectively generalize their map knowledge beyond their actual experiences, for example, filling in gaps between adjacent locations or reversing their direction. This is similar to other models that permit the application of multiple episodic memories to improve generalization (Rikhye et al., 2019).

#### Nonparametric model

So far, we discussed a generative model with a fixed number of clusters (five). Here, we illustrate the same continual learning experiment using a nonparametric extension of the model, which permits an open-ended expansion of the number of clusters as the agent receives novel observations (see the Methods section for details). This setup more closely reflects the problems that the hippocampus has to solve, given that in realistic settings, the number of clusters is not known a priori. Unlike the previous simulations, here each maze was presented 4 times each, across 20 navigation blocks. This was done because nonparametric methods generally require more training data than parametric methods, as they need to infer the appropriate number of clusters. We trained 16 replicas of each agent.

The performance of the nonparametric method approximated the parametric method both during learning and retention; compare Figures 4C-D and 1C-D. During learning, the agents with generative replay and prioritized generative replay outperformed the baseline agent (t=2.47, s.d.=0.10, df=30, p=0.02 and t=4.04, s.d.=0.09, df=30, p<0.001, respectively) and – in contrast to the experiment using the parametric approach – also the experience replay agent (t=1.13, s.d.=0.09, df=30, n.s. and t=2.67, s.d.=0.08, df=30, p=0.01, respectively). The analysis of the retention test showed an even stronger advantage of the generative approach: the generative replay and prioritized generative replay agents outperformed the baseline agent (t=2.89, s.d.=0.09, df=30, p=0.007 and t=4.18, s.d.=0.09, df=30, p<0.001, respectively) and the experience replay agent (t=2.76, s.d.=0.07, df=30, p=0.01 and t=4.68, s.d.=0.06, df=30, p<0.001, respectively). The two agents using generative replay and prioritized generative replay showed no significant differences in either learning (t=1.437, n.s.) or retention (t=1.487, n.s.).

**Figure 4.**
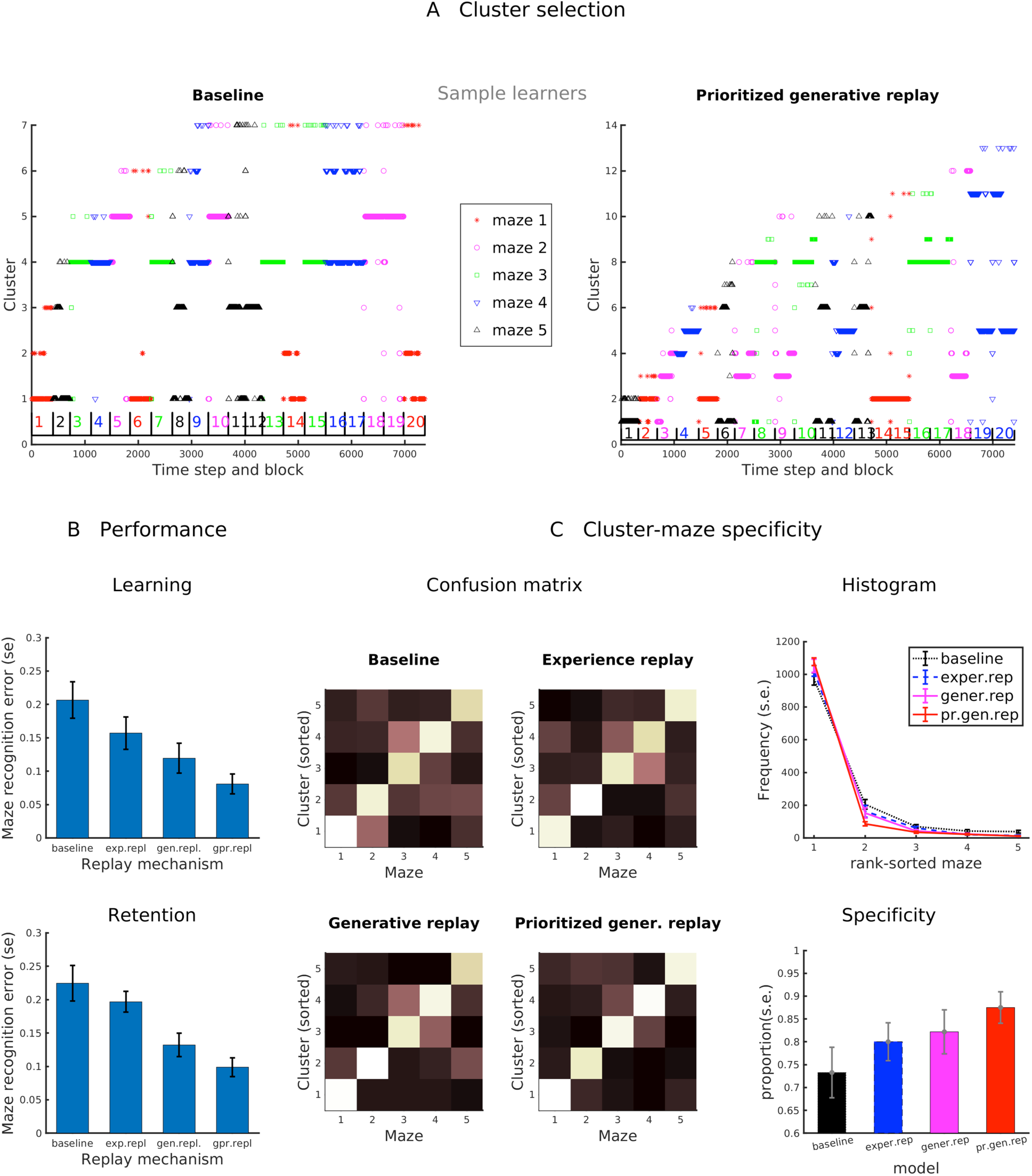
Clustering emerged during continuous nonparametric learning. (A) Cluster selection during navigation in sample baseline and prioritized generative replay agents. Note the greater maze-cluster consistency in the latter agent. (B) Performance of the four models during learning and retention (as in Fig 1C,D). (C) Analysis of the consistency of cluster-maze selection across all replicas of the four learning agents. Left: Average cluster-maze confusion of the clusters that are most frequently selected during the navigation in each maze (note that this analysis regards in total n=5 clusters per learner). Right: as in Figure 2, average frequency of cluster selection and proportion of top maze-specific clusters selected.

As expected from a nonparametric method, all the agents showed some redundancy in their learned clusters (Figure 4A) but were still able to reliably map each maze to at least one cluster – as evident in the confusion matrices in Figure 4C. Similar to our parametric results, the two agents using generative replay and prioritized generative replay developed more specific clusters, which may explain their improved performance. These results illustrate that our approach to continual learning does not necessarily require a-priori information about the number of clusters.

#### Comparison of the hierarchical model with reduced models that lack hierarchical depth

So far, we have discussed a hierarchical generative model that comprises three layers, for items, sequences and maps, respectively (Figure 1). Here, we test the usefulness of this hierarchical structure, by comparing the performance of agents that use the full hierarchical (parametric) model with two reduced models that lack hierarchical depth. The first reduced model is the same as the full model, but lacks the sequence layer. The second reduced model is the same as the full model, but lacks the ability to form different maps, because it only includes 1 cluster (instead of 5) at the highest level.

We compared the same four agents (baseline, experience replay, generative replay, prioritized generative replay) using the full hierarchical model with equivalent agents that used the two reduced models, in four tests. The first two tests were the same Learning and Retention tests discussed above (see Figure 1C-D for the results of the four agents using the full hierarchical model). We found that if the same four agents use the first reduced model that lacks the sequence layer, their performance in both the Learning and Retention tests dropped significantly (all p<0.001; see Figure S1). Furthermore, the four agents developed poorly differentiated spatial maps at the highest level (see Figure S2). This result shows that including the sequence layer, which provides a coherent (temporal) context to organize individual items of experience, is crucial to learning effective spatial maps. Note that the second reduced model that we considered cannot by definition solve the Learning and Retention tasks, as it includes only one cluster at the highest (map) layer and hence cannot differentiate between mazes. Unsurprisingly, we found that the second reduced model learns a map that combines all the experienced mazes (see Figure S3).

Our third test considers how many impossible locations (i.e., those that violate maze constraints) the four agents replay, when using the full model or the two reduced models (see Figure S4A-C). Our results illustrate that when averaging the performance of the four agents, the full model generates significantly fewer impossible locations compared to the reduced model that lacks the sequence layer (p=0.000, t=-15.498, df=30, sd=0.012) and to the reduced model that only includes 1 cluster (p=0.000, t=-15.271, df=30, sd=0.014).

For completeness, we tested the differences between the four agents that use the full model. We found that the baseline agent replays significantly more impossible locations compared to the experience replay agent (p=0.023, t=2.404, df=30 sd=0.014), the generative replay agent (p=0.032, t=2.244, df=30, sd=0.014) and the prioritized replay agent (p=0.016, t=2.547, df=30, sd=0.014). This finding complements our previous results, showing that using replay during learning leads to better generative models that do not violate environmental constraints.

Finally, our fourth test considers the confidence of the agents’ predictions during replay, defined here as the peak probability of each predicted location (see Figure 5D-F). Using this definition, the higher the probability assigned to the next predicted location, the higher the agent’s confidence. This test is compelling to the extent that generating predictions with high confidence may index a more robust model (to the extent that the predictions are also largely correct, as assessed above). Our results illustrate that when averaging the performance of the four agents, the full model generates significantly fewer impossible locations compared to the reduced model that lacks the sequence layer (p=0.001, t=3.612, df=30, sd=0.007) and to the reduced model that only includes 1 cluster (p=0.000, t=34.661, df=30, sd=0.005).

**Figure 5.**
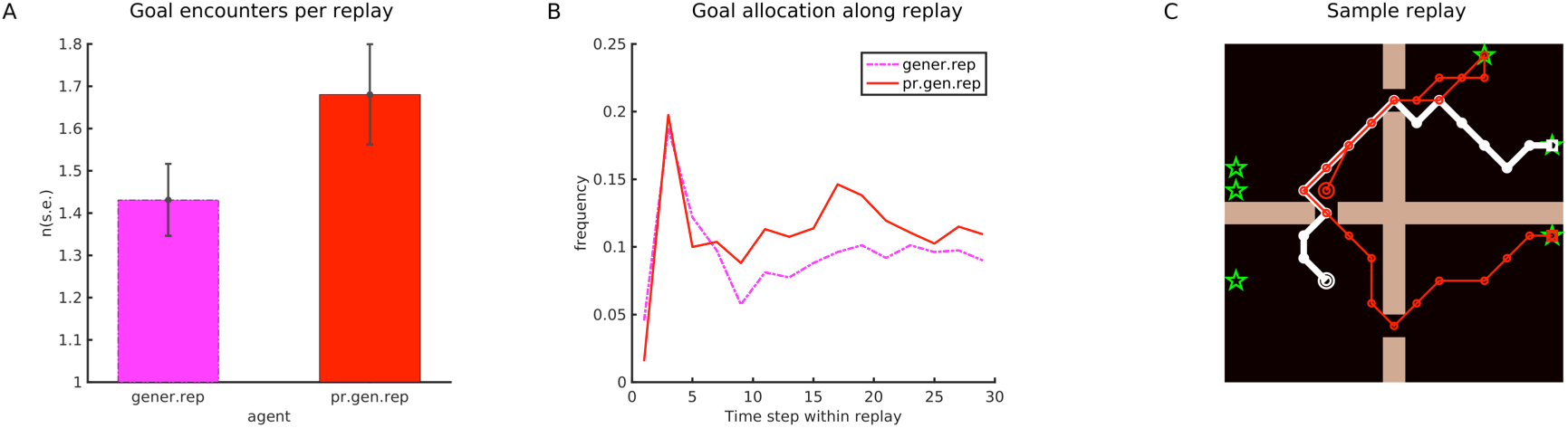
Analysis of goal-sensitivity during generative replays in the absence of external stimuli (akin to hippocampal replays during sleep). (A) Frequency of fictive visits to goal locations during generative replay. (B) Frequency of fictive visits to goal locations during the generative replays (lasting 30 time steps for simplicity). Generative replays tend to start close to goal locations, akin to what shown empirically; but show also goal sensitivity during the whole fictive trajectories. (C) An example novel trajectory (red) produced during replay that shares some parts with an experienced trajectory (white). The novel trajectory starts from the double circle in the top-left quadrant, then reaches two goal locations (green stars), in the top-right and bottom-right quadrants, respectively.

We also found that in the full model, the experience replay (p=0.009, t=-2.774, df=30, sd=0.010), generative replay (p=0.000, t=-11.012, df=30, sd=0.011) and prioritized generative replay (p=0.000, t=-14.053, df=30, sd=0.010) agents all generate higher confidence predictions compared to the baseline model. Furthermore, both the generative replay (p=0.000, t=-9.472, df=30, sd=0.010) and the prioritized generative replay (p=0.000, t=- 12.847, df=30, sd=0.009) agents generate higher confidence predictions compared to the experience replay agents, suggesting that generative replay might lead to more robust models.

Taken together, the results of the continual learning experiment shows that agents endowed with a hierarchical generative model and generative replay mechanisms can efficiently encode multiple mazes, infer its current maze, and generate realistic fictive experiences. Furthermore, the hierarchical generative replay model proposed here consistently outperforms models that lack replay, models that use experience replay (especially in the more compelling nonparametric setting) and models that lack hierarchical depth. Taken together, these results support the computational efficiency of the novel proposed model. In the next section, we move from this computational-level analysis to a biological-level analysis, by discussing the possible relations between the model’s inferential dynamics and internally generated hippocampal sequences.

### Neurobiological underpinnings of generative replay

We have argued that generative replay provides a novel perspective on internally generated hippocampal sequences, suggesting that they may be better understood as *sequences resampled from a latent model*, rather than the verbatim replay of previous experiences as commonly assumed. Here, we explain some empirical implications of this novel perspective.

#### Generative replay in different modalities: during sleep and wakeful replay

In rodents and humans, replays have been observed in two main modalities: during sleep and during wakeful rest, such as when an animal pauses between navigational experiences (Foster, 2017). Here we propose that replays during both sleep and during wakeful rest can be conceptualized as generative replay, with different contextual conditions: when stimuli are largely absent and when they are present, respectively.

When external stimuli are absent or weak (e.g., during sleep), generative replay produces sequences entirely composed of fictive observations; this mechanism is sometimes called an *inputless decoder* in machine learning. Importantly, during learning, the generative agents replay sequences from multiple maps, not just from the most recently experienced one (they select a map for each replay, based on current cluster probabilities). This mechanism nicely explains why hippocampal replays can include distal experiences rather than just the last visited (portion of the) maze (Gupta et al., 2010). Furthermore, the generative episodes start from an item / location *sampled from* one of the maps. This is important because the maps encode the probability of items at given locations (i.e., empirical priors in Bayesian terms), reflecting their behavioural relevance. Hence, in the absence of external stimuli, generative replay events will tend to start from behaviourally relevant locations, such as goal locations and frequently visited junction points of the problem (see Figure 5). This result aligns well with the finding that hippocampal replay during sleep tends to start form rewarded locations and proceed “backwards” to the trial’s start position (Ambrose et al., 2016). The same goal sensitivity emerges in prioritized generative replay agents, despite generating replays based on maps that encode (Shannon) *surprise*, rather than probabilities. This is because an encounter with novel goals or rewards is registered as surprising in the maps used by prioritized generative replay agents. Crucially, the selective resampling of goal (and other behaviourally relevant) items or locations further “biases” the maps to over-represent these items or locations, above and beyond actual experience. This result is in keeping with previous formal analyses suggesting that (backward) replay could help learning goal-directed reinforcement learning policies (Ambrose et al., 2016; Mattar and Daw, 2017).

Generative replay can also occur when the agent momentarily disengages from a behavioural task, akin to *wakeful replay* in rodents. In this case, the agent receives richer bottom-up inputs, such as observations about its current location, which can provide context and a starting point for generative replay. Generative replay would therefore start from the (currently inferred) location and produce sequences of fictive observations, which proceed along a path of high probability in the (currently inferred) map. As the high-probability paths in the maps typically encode behaviourally relevant trajectories, generative replay could potentially support goal-directed planning and the optimization of the model for the current task. This aligns well with the finding that compared to replay during sleep, replay during wakefulness is more likely to start around the animal’s position and to proceed towards behaviourally relevant locations (Pfeiffer and Foster, 2013; Widloski and Foster, 2022).

In sum, our model suggests that replays during both sleep and wakefulness (and potentially also theta sequences, see the discussion) may be manifestations of the dynamics of the same hierarchical generative model, operating in different conditions, with or without strong bottom-up inputs to initialize a generative episode.

#### Replay statistics: similarity of spontaneous activity before and after navigation

A recent study showed that hippocampal replay statistics *after* navigation in a novel maze are better explained by sequential dynamics that already existed in the hippocampus *before* navigation, than by recent navigational experience (K. Liu et al., 2019). Specifically, the study reported rank-order correlations between pre-run hippocampal activity (i.e., replays measured before navigational experience) and post-run activity (i.e., replays measured after navigational experience) to be significantly greater than rank-order correlations between run activity (i.e., hippocampal activity measured during navigational experience) and post-run activity. This finding suggests that hippocampal temporal coding depends primarily on selective changes to preconfigured network dynamics (Dragoi, 2020).

To test whether our model reproduces the above empirical findings, we performed the same rank-order analysis as in (K. Liu et al., 2019) on our generative replay and prioritized generative replay agents. We firstly calculated temporal rank-order maps (rank-maps, for short) of all *observation* units shown in Figure 1 before navigational experience (pre-run), during navigational experience (run) and after navigational experience (post-run) for each of the five experimental blocks. We then measured the correlation between pairs of rank maps: *post-run* with *pre-run*, and *run* with *post-run*.

Please note that the *observations* used for the rank maps correspond to locations that are either actually observed during navigational experience (*run*) or spontaneously predicted before (*pre-run*) or after (*post-run*) navigational experience. For each trajectory, the observation-level units participating in the trajectory were ranked according to their temporal order of activation in the trajectory. Figure 6A shows the rank-maps of a sample agent during a navigation episode, with the colour of each cell representing the rank of the observation in the corresponding location, from black (lowest) to white (highest) rank.

**Figure 6.**
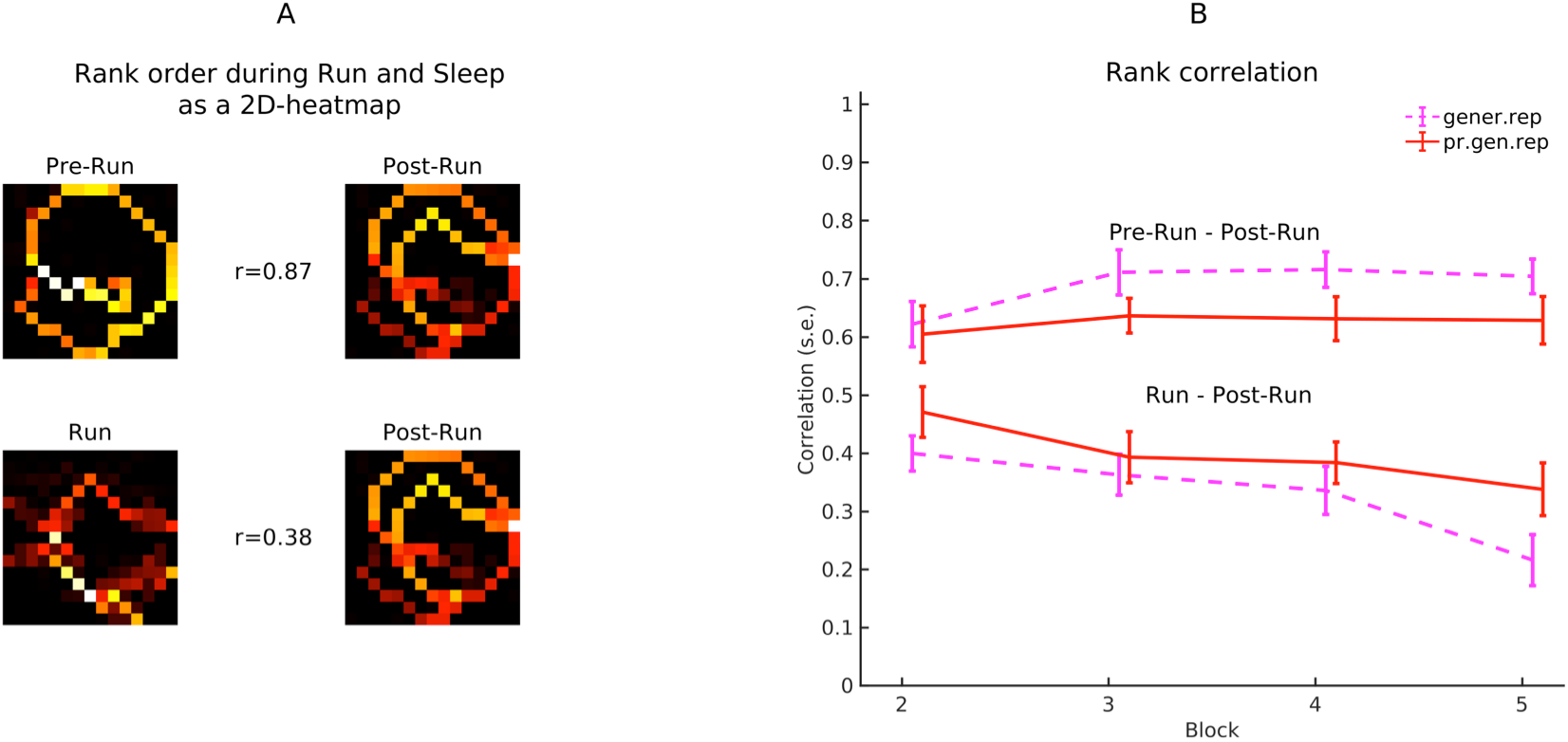
Network dynamics before novel navigation experiences explain replay activity better than novel experiences themselves. (A) Rank order maps of a sample learner during sample pre-run, run and post-run phases, along with rank correlations between pre-run and post-run, and run and post-run, respectively. Colours represent rank, from white (high) to black (zero). The ranks were normalized to 1 and the normalized ranks of all trajectories from a given episode were accumulated in a 2D rank-map, which was therefore normalized to sum 1. (B) Rank-correlation statistics across all n=16 replicas for each of the n=2 conditions (generative replay and prioritized generative replay) and n=5 learning blocks.

Spearman’s rank order correlation was used to compute the correlations between the post-run and pre-run rank maps and between run and post-run rank maps, separately for each learning condition, replica, and block. Figure 6B shows statistical plots for each condition and episode across all 16 replicas per condition. The post-run – pre-run correlations were compared with the post-run – run correlations using two-sample t-tests and effects of block were explored using a linear regression of the correlations with block and condition as predictors, i.e. Rank-Correlation ~ condition + block.

Consistent with previous findings (K. Liu et al., 2019), the average pre-run – post-run rank correlation across conditions, blocks, and replicas (0.66, s.d.=0.15, n=128) was significantly greater than the average post-run – run correlation (0.36, s.d.=0.17, n=127; t=17.05, p<0.001). Note that while the original rodent study (K. Liu et al., 2019) focused on a single maze, here we calculated correlations across multiple, sequentially learned mazes (in pseudorandom order).

The reason why the trajectories spontaneously generated by the clusters before navigational experience resemble those generated after it is that the clusters are not selected randomly to encode one map or the other. Rather, when learning a particular maze, the model selects the cluster that already expresses the spontaneous repertoire that fits the maze better. This means that the map with the best-preconfigured spontaneous repertoire is selected and later adapted to encode each particular map. Interestingly, we also found that the run – post-run rank correlations steadily decreased across blocks in a linear regression fit (β=-0.05, t=-3.9, p<0.001), which was not the case for the pre-run – post-run correlations (β=0.02, t= 1.35, p=0.17). This constitutes a novel prediction of our framework, which remains to be tested empirically.

#### Replay statistics: spatial diffusion of replays

A recent study reported that replays followed Brownian diffusion-like random trajectories even if the animal’s trajectories during navigation were not Brownian (Stella et al., 2019). Brownian diffusion characterizes the average Euclidean distance Δ*x* between any two locations along the path of a randomly moving particle as a power law function of travel time Δ*t*, i.e., Δ*x* ≈ Δ*t^α^* where *α* is about 0.5. This can be statistically quantified by calculating the regression slope α between the average log-Euclidian distance in space covered by neural trajectories and the corresponding log-travel time. In the animal data it was found that α ranges between 0.45-0.53 during replay (Figure 3A of (Stella et al., 2019)) and about 0.67-0.72 during navigation (Figure 4A of (Stella et al., 2019)) demonstrating that that the hippocampus can generate trajectories with statistics different from those experienced during navigation. This is similar to what happens in our model, where generative replay produces novel trajectories that are compatible with the learned maps.

We performed the same log-log regression analysis on the trajectories of our generative replay and prioritized generative replay models (considered together); see Figure 7A. Consistent with empirical findings, our model generates trajectories which approximate Brownian motion (average log-log regression coefficient across all learners and conditions α=0.46, s.d.=0.031, n=32), despite trajectories experienced during navigation reaching greater distances for the same time period (average regression coefficient α=0.75, s.d.=0.014). Figure 7B provides an intuitive illustration of these results, by showing the relative spatial diffusion of navigation and replay trajectories starting from either a central zone or the periphery (we considered trajectories starting at the 3rd time step to ensure that maps were correctly inferred). To compare the diffusion maps during navigation and replay, we calculated the average entropy of the relative diffusion maps across 5 time steps and 2 conditions (centre and periphery). Entropy was higher during replay (lower 2 rows: H = 6.63, s.d. 0.69, n=10) than during navigation (upper two rows: H = 4.89, s.d. 1.21, n=10). Overall, the quantitative and visual analysis showed that diffusion, as in animal studies, was much more directed during navigation than during replay.

**Figure 7.**
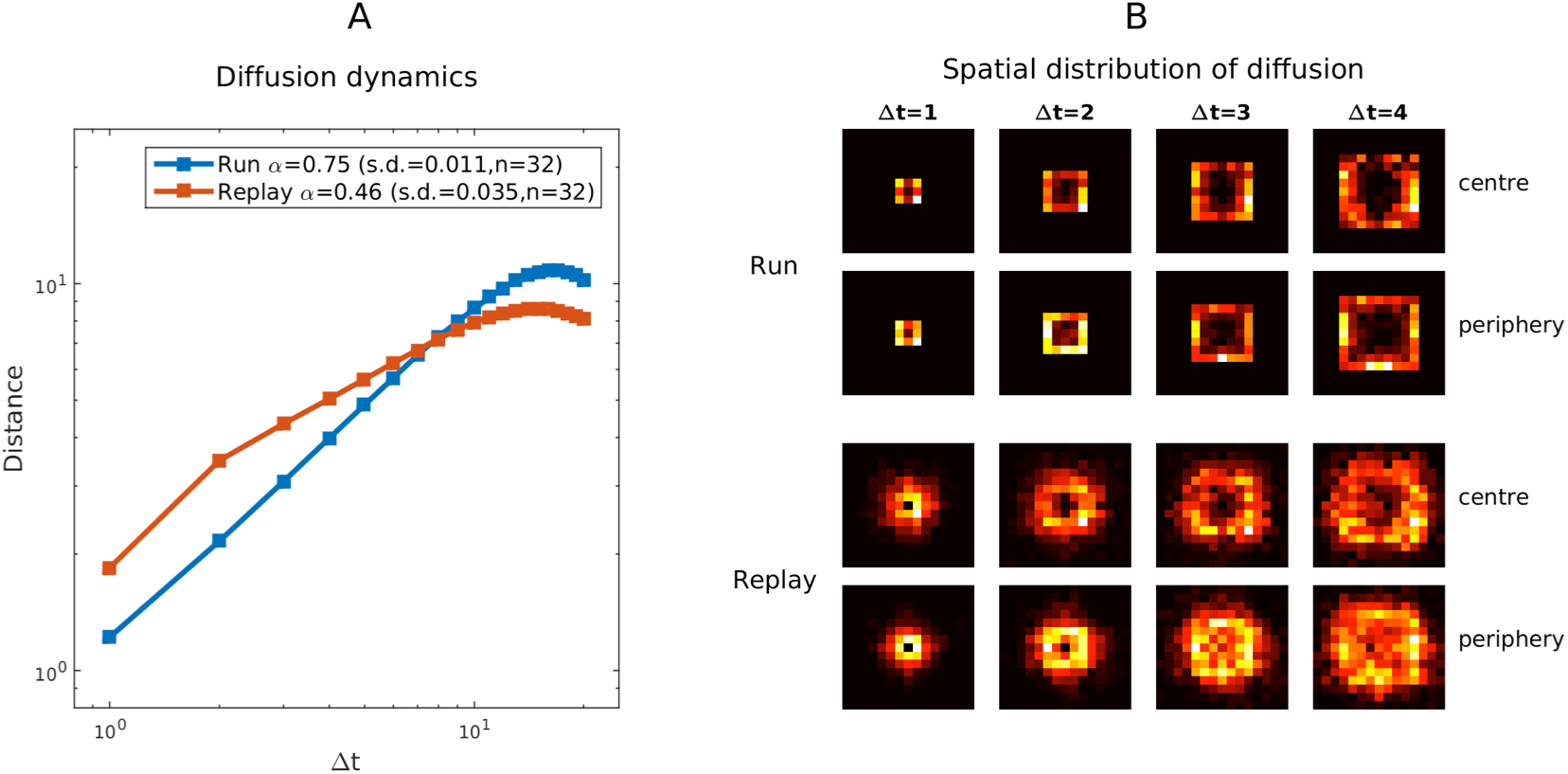
Trajectories during spatial experience and during replay across all generative replay learners and conditions. (A) Average point-to-point distance as a function of time-step intervals, plotted on a log-log scale. (B) Relative diffusion of locations during navigation and replay at five time steps for trajectories originating from any point of the central zone, i.e., the 5×5 central square (first and third row of panels) or the periphery, i.e., the remaining locations (second and fourth row of panels).

#### Hippocampal coding of cognitive maps

Our model assumes that the hippocampus infers cognitive maps: structured representations that define a coherent context to organize stimuli, forming spatiotemporal codes for spatial navigation (Plitt and Giocomo, 2019), but also possibly non-spatial codes for other structured tasks (Aronov et al., 2017; Constantinescu et al., 2016; Gulli et al., 2020; Mok and Love, 2019). While our analyses focused mostly on a parametric model that uses a fixed number of clusters, the nonparametric extension explored in the section “Nonparametric model” is better suited to capture the ability of the hippocampus to encode a very large number of experiences and to restructure itself during development and learning.

In the model, relational and contextual information is present at the level of sequences and maps, but not at the level of individual item codes. This is in keeping with evidence that in a structured, multi-compartment maze, place cells code for spatial locations but not connectivity (Duvelle et al., 2021). It is important to note that while item codes of different maps are visualized in the same positions of the 15×15 grids of Figure 1, they may actually correspond to the activation of different hippocampal neurons. In other words, different maps can be neurally encoded using different sets of hippocampal neurons, or by partially overlapping ensembles of neurons that “remap” across multiple maps – e.g., two neurons coding for adjacent locations in one map may code for non-adjacent locations in another map. It is this remapping mechanism that enables the disambiguation between two identical trajectories executed in two distinct spatial environments; and more broadly, avoids interference between similar experiences (Colgin et al., 2008).

Our simulation results show that with enough capacity, the model tends to group multiple coherent experiences in a single cluster or map – which is useful for generalization and to generate novel, never experienced trajectories; see Figure 2B. However, the model does not necessarily acquire a one-to-one correspondence between clusters and mazes. Depending on the training regime and the similarity between maps, the model can in some cases develop multiple clusters for the same maze, or reuse the same clusters across multiple mazes. The first possibility – using multiple clusters for the same maze – illustrates a potential mechanism for hippocampal splitter cells (Grieves et al., 2016b), which show route-dependent rather than map-dependent firing. The second possibility – reusing the same cluster across multiple mazes – may explain repeating place fields often observed in repetitive (e.g., multicompartment) environments (Grieves et al., 2016a; Spiers et al., 2015). Furthermore, this dynamic provides a potential mechanism for generating never-observed sequences that cross two mazes, or two portions of the same maze that were never experienced together, e.g., shortcut sequences (Gupta et al., 2010); see also (Y. Liu et al., 2018). These shortcut sequences were empirically found while animals independently learned two portions of the same maze – where distal cues (not included in our model) may have facilitated the integration of these memories and avoided segregation into distinct maps (Gupta et al., 2010). More broadly, reusing the same clusters across multiple mazes provides an effective mechanism for the rapid learning of novel spatial maps from a very limited set of experiences – or even prior to navigation experience (Dragoi and Tonegawa, 2011).

A key distinguishing feature of this model is the use of a mixture model for map probabilities, which implies that only one map is selected at a time during learning and inference. While this may be a simplification, two key phenomena – theta-paced flickering (Jezek et al., 2011) and the constant cycling between representations of different trajectories during navigation (Kay et al., 2020) – suggest that place cell assemblies from different mazes are activated within different theta cycles, and map representations are not combined. To assess whether and in which conditions flickering occurs in our model, we performed an analysis of the time-course of the maze inference performed by the four agents during a variant of the Retention test, in which we did not reset the cluster probabilities after each trial. In this task, the agents had to infer the correct maze for two consecutive trials, in which the maze remained the same (“same” condition) or changed (“change” condition), as if they were “teleported” to a novel maze (Jezek et al., 2011). To make the “change” condition more challenging, we only considered changes between the mazes that are most similar (i.e., 1-2 or 3-4). We considered whether and how often flickering (i.e., changes from correct to incorrect map inference, before the inference settles) occurred in the second trials of the test, see Supplementary Figure S7. Figure S7a shows some example trials in the “change” condition that include flickering, for four sample learners. Here, flickering corresponds to a sequence formed by a white square (i.e., correct inference) and then one or more black squares (i.e., incorrect inference). As the figure shows, flickering occurs across all the agents, sometimes multiple times per trial. Figure S7b shows the statistics of flickering in the “same” and “change” conditions, for 16 replicas of each agent. The figure shows that flickering is more common at the beginning of “change” trials, similar to what is reported in Figure 3e of (Jezek et al., 2011). Finally, Figure S7c shows that across all the agents, most flickering events have a short duration (i.e., 1-4 inferential steps), similar to what is reported in Figure 3f of (Jezek et al., 2011). This control simulation shows that flickering is a consequence of the hierarchical architecture as it occurs (with a different frequency) across all the agents and it is more prominent after a “teleport”, as reported in (Jezek et al., 2011).

#### Putative biological implementation of generative modelling in the hippocampal formation

We argued that the hippocampal formation functions as a hierarchical generative model that organizes sequential experiences into coherent but mutually exclusive spatiotemporal contexts – or cognitive maps. The most widely known neurobiological theory of generative modelling and inference is predictive coding (Friston, 2005; Rao and Ballard, 1999). Predictive coding is based on a hierarchical brain architecture with reciprocal feedfoward and feedback connections, which is common in the cortex. However, the extended hippocampal formation follows a different anatomical scheme, in which (with some simplifications) the entorhinal cortext and the hippocampus form a large recurrent loop (Kloosterman et al., 2004; Koster et al., 2018): the entorhinal cortex provides “inputs” to the hippocampus (via the dentate gyrus and CA1) and in turn receives its “outputs” (via CA1 and the subiculum), recirculating them.

A speculative possibility is that this recurrent architecture might support both inference (or *encoding*) and generation (or *decoding*) functions of generative models, in distinct pathways. The inference of items, sequences and maps could proceed from the lateral and medial portions of the entorhinal cortex (which provide sensory and spatial information, respectively) through the dentate gyrus (which forms pattern-separated, conjunctive item codes, such as place cells) and culminate in CA3 and/or CA1. Conversely, decoding could proceed from CA3 to CA1 and back to the entorhinal cortex. The recurrent architecture could permit iterating phases of inference and generation, as required in our model when (for example) the model infers the current map and then generates a replay from this map.

In this scheme, temporal and relational-spatial codes of the hierarchical generative model might map to hippocampal and entorhinal structures, respectively. In other words, the hippocampus could encode sequential episodes (corresponding in our model to item and sequence codes) and the entorhinal cortex could provide a metric structure to integrate experiences (corresponding in our model to map codes and the “2D squares” shown in Figure 1A) or encode (average) transition models (Evans and Burgess, 2019; Whittington et al., 2018). An alternative possibility is that the three levels of the generative model are not localized in distinct areas, but rather distributed across the extended hippocampal formation, with both hippocampal (place) and entorhinal (grid) codes operating as “basis functions” to encode maps (Whittington et al., 2020). This distributed scheme might account more easily for the fact that the entorhinal cortex codes not just spatial or topological information but also reward and goal information (Boccara et al., 2019; Butler et al., 2019). Whether temporal and relational-spatial codes of the generative model are localized or distributed across the hippocampal formation remains to be fully assessed empirically.

Other brain areas outside the extended hippocampal formation may be related to our proposed generative model. While internally generated hippocampal dynamics are largely dependent on intrinsic (CA3) dynamics, they are usually coordinated with other brain structures, such as medial entorhinal cortex (Chenani et al., 2019) and mPFC (Shin et al., 2019; Tang et al., 2020). Prefrontal structures may play a facilitating role for hippocampal sequences to occur when needed, for example during decision-making (Redish, 2016), and hence mediate prospective (planning) and retrospective (learning) functions of hippocampal replay (Foster, 2017; Penagos et al., 2017; Giovanni Pezzulo et al., 2017; Pezzulo et al., 2019). Furthermore, coordinated sequential replay across the hippocampus and PFC (Tang et al., 2020) and hippocampus and ventral striatum (Lansink et al., 2009) may support goal-directed spatial navigation (Pezzulo et al., 2014; van der Meer et al., 2012). The information exchange between hippocampal and cortical structures may also support bidirectional learning. It has long been proposed that the hippocampal model can train cortical semantic systems or behavioural controllers offline. In turn, cortical systems can transfer structured, rule-related knowledge to hippocampal systems (Penagos et al., 2017), potentially biasing their content and reorganizing experiences according to learned rules (Y. Liu et al., 2019). A theoretical possibility that remains to be tested in future research is that hippocampal and cortical structures both implement generative models that replay simultaneously and train each other during periods of coupled oscillations.

## Discussion

We have proposed a novel theory of the hippocampal formation as a hierarchical generative model of spatiotemporal experiences. The theory proposes that the hippocampal generative model is hierarchically organized and includes three layers of hidden states: *items* (that disentangle representations), *sequences* (that organize items in time), and *maps* (that organize sequences in space). A key role of the hippocampal generative model is therefore organizing individual items of experience coherent spatiotemporal contexts, allowing the model to link multiple observations experienced across different navigational episodes (Gupta et al., 2010). This idea is compatible with the well-established roles of the hippocampus in cognitive map formation (Tolman, 1948), relational tasks (Eichenbaum, 2000) and the spontaneous generation of sequential dynamics (Buzsáki and Tingley, 2018) making it a quintessential “sequence generator” (Buzsáki and Tingley, 2018; Giovanni Pezzulo et al., 2017).

Furthermore, the theory suggests that the hippocampus maintains several independent maps that correspond to different relational contexts in which experiences were made. From a computational perspective, the capability to learn and discriminate different contexts is key to learn multiple experiences while avoiding catastrophic forgetting (Rikhye et al., 2020); whereas from a biological perspective, it might be key to support episodic and autobiographical memory (Tulving, 2002). Importantly, in our model, map learning starts with the selection between multiple preconfigured maps, which might sometimes correspond to different sets of neurons (Jezek et al., 2011). This favours the selection of maps that are already preconfigured for novel mazes, hence explaining why network dynamics before novel navigation experiences explain replay activity better than novel experiences themselves (K. Liu et al., 2019), see Figure 6. This approach has some resemblances with “selectionist” theories of learning (Edelman, 1987), with machine learning approaches such as *reservoir computing* that select between a preconfigured set of dynamics (Dominey et al., 1995; Jaeger et al., 2007; Lukoševičius and Jaeger, 2009) and the “lottery ticket” hypothesis that learning in large neural networks is largely driven by subnetworks having fortuitous initializations that render them appropriate for the task at hand (Frankle and Carbin, 2019).

Finally, the theory proposes that the hippocampal generative model supports *generative replay*: a method for continual learning that uses fictive sequences generated from a learned model to train the model itself, or other (e.g., cortical) models or controllers. Accordingly, spontaneous hippocampal dynamics are manifestations of generative replay. Our proposal is related to the widespread idea that the hippocampus uses experience replay to improve learning (Kumaran et al., 2016; McClelland et al., 1995; Mnih et al., 2015), but differs from it in fundamental ways. First, the above proposals stress the importance of replay to train cortical models outside the hippocampus, whereas generative replay serves first and foremost to train the hippocampal generative model itself (but can train other systems in parallel). Second, experience replay as used in machine learning is not biologically feasible, as it requires a system capable of storing an unbounded number of perfect memory traces. The architecture proposed here exemplifies a possible way the brain might do something like experience replay, but without perfect and unbounded memory. An alternative possibility is that the brain could replay from an auto-associative memory network that stores static memory patterns (Treves and Rolls, 1992) or sequences of experiences (Cheng, 2013; Levy, 1996; Lisman, 1999). While these possibilities seem similar, replaying from a generative model of experiences (as we propose) is different than replaying from a noisy memory of the experiences (as would be the case with an auto-associative memory network). Several empirical studies indicate that the statistics of hippocampal replay events are significantly different from a noisy recapitulation of experiences and seem to be better captured by appealing to generative replay, see the section on “Neurobiological underpinnings of generative replay”. Finally, in our proposed model, generative replay largely exploits intrinsic network dynamics - here, partially preconfigured maps - rather than just navigational experiences. This is another commonly reported finding that is difficult to reconcile with the idea that replays come from noisy memories (K. Liu et al., 2019).

Our proposal is appealing from both machine learning and biological perspectives. From a machine learning perspective, generative replay prevents catastrophic forgetting (Shin et al., 2017b) without requiring the verbatim storage of an unbounded number of previous experiences, as in “experience replay”. Our results show that generative replay and prioritized replay agents outperform baseline agents (that do not use replay) in all our tests and also agents using experience replay (i.e., replaying from a memory buffer) in the most challenging, nonparametric setting. Across a series of simulations, we were able to assess that what makes the model effective for continual learning is the combination of hierarchical structure and generative replay. The hierarchical structure permits organizing experiences into multiple maps, providing resistance to catastrophic forgetting. Generative replay permits learning the hierarchical models better than without replay and at the same level of accuracy of (or even better than) ideal experience replay, which is biologically unachievable.

At the biological level, generative replay explains the flexibility of internally generated hippocampal sequences, which are difficult to reconcile with previous theories that consider them rigid “replays” of previous experiences. This is supported by three observations. First, the hippocampus can generate novel sequences that show prospective aspects. These are explained in our model by considering that sequences follow high probability paths, and the maps from which sequences are generated implicitly encode behaviourally relevant or salient locations (Ambrose et al., 2016; Bhattarai et al., 2020; Igata et al., 2021; Ólafsdóttir et al., 2015; Pfeiffer and Foster, 2013; Wikenheiser and Redish, 2015). From a conceptual perspective, while sequences encode short-term predictions, maps encode long-term predictions of future locations that average across several episodes; this renders our idea compatible with the view that the hippocampus learns a predictive representation (Recanatesi et al., 2018) but casts it within a hierarchical inference scheme.

Furthermore, in our model maps can express a rich repertoire of sequential dynamics even before a specific navigational episode. The map that best fits the novel navigational episode is then selected and adapted to incorporate the experience. This mechanism implies that sequences generated before a navigational episode – preplays (Dragoi and Tonegawa, 2011) – and after it – replays – are highly correlated, as demonstrated empirically (K. Liu et al., 2019). In turn, this points to the importance of preconfigured sequential dynamics to explain fast learning in the hippocampus (Dragoi and Tonegawa, 2013; K. Liu et al., 2018).

Finally, in keeping with empirical findings (Stella et al., 2019), the sequences produced during generative replay follow Brownian, diffusion-like dynamics, while navigational experiences do not. This finding further exemplifies the generative nature of hippocampal sequences, which do not simply recapitulate recent experiences, either in terms of content or dynamics.

It remains to be fully assessed which of our generative replay models (or perhaps a combination of both) best explains the phenomenology of hippocampal sequences. A previous formal theory considered replays to be guided by two terms: a *need* term that prioritizes behaviourally relevant experience and a *gain* term that prioritizes novel experiences; but it did not provide a mechanistic implementation of this idea (Mattar and Daw, 2017). By noticing that need and gain terms have close correspondences with the functioning of generative replay and prioritized generative replay, respectively, our proposal could shed light on the potential mechanisms supporting these two factors that contribute to explain replay. Another possibility that remains to be tested is that the two generative replay mechanisms may be selected in context-dependent ways, depending on the activity of different neuromodulators, which are known to influence replay activity (Atherton et al., 2015).

### Relations with other proposals

Previous theoretical and computational studies have assigned a role to the hippocampus in supporting the encoding of sequential memories and model-based, goal-directed navigation (Cazin et al., 2019; Cheng, 2013; Lisman and Redish, 2009, 2009; Penagos et al., 2017; Penny et al., 2013; Pezzulo et al., 2012; Giovanni Pezzulo et al., 2017; Pezzulo et al., 2021; Stoianov et al., 2018; Wittkuhn et al., 2021). Our proposal is consistent with these ideas but goes further by introducing the idea of hierarchical, structured representations of sequences and maps, and by explicitly modelling how these representations are learned over time.

Other studies used a probabilistic approach to simulate and analyse replays. (Evans and Burgess, 2019) advanced a probabilistic model of structural inference and learning in the hippocampal-entorhinal network and used it to simulate a coordinated replay across the two structures. Furthermore, they demonstrated how Hebbian learning in place and grid cells may support the proposed inferential scheme in a biologically plausible way. Using a similar generative approach, (McNamee et al., 2021) simulated replay and other forms of hippocampal reactivations in a linear feedback model of the hippocampus and entorhinal cortex. A specific parameterization of the model includes spatial inductive biases that are largely equivalent to those used in this manuscript (i.e., space is smooth and isotropic). Furthermore, the model shows that modulating the grid population input permits interpolating between distinct regimes of sequential hippocampal reactivations; namely, the diffusive reactivations observed during sleep (Stella et al., 2019) and simulated in Figure 7 of this paper, superdiffusive patterns observed in the awake behaving state (Krause and Drugowitsch, 2022; Pfeiffer and Foster, 2015, 2013) and minimally autocorrelated sampling that resembles generative cycling during navigation (Kay et al., 2020). Finally, (Krause and Drugowitsch, 2022) used a probabilistic generative model to simulate hippocampal reactivations having the same spatial priors as in this manuscript and (McNamee et al., 2021), but also included momentum, which permits trajectories to be ballistic. While all these computational models are based on an inferential approach and share similarities with our proposal, they do not use a hierarchical representation and hence might be less suited for the continual learning of multiple mazes, similar to our control simulations with reduced models that lack hierarchical depth.

The generative replay mechanism investigated here shares resemblances with the DYNA architecture of reinforcement learning (Sutton, 1991), which self-generates experiences from a learned *transition model* of the task, to train a reactive reinforcement learning controller offline. This idea has inspired the recent proposal that spontaneous hippocampal activity might serve to optimize reinforcement learning controllers (Mattar and Daw, 2017). There are two main differences between our proposal and these alternative theories. First, our proposal views the hippocampal formation as a hierarchical generative model of spatiotemporal experiences, not a transition model. The former (generative model) denotes a statistical model of the joint probability distribution on observations and hidden variables (here, map, sequence and item variables, but not action variables). The latter (transition model) denotes the conditional probability distribution of future states given the current state and action. Second, our proposal emphasizes the importance of spontaneous activity to improve the generative model and optimize it for future use (Giovanni Pezzulo et al., 2017; Pezzulo et al., 2021), rather than to learn reactive reinforcement learning controllers, as in DYNA (Sutton, 1991) and related proposals (Mattar and Daw, 2017). From a computational perspective, learning a generative model (as done in generative replay) is simpler and less error prone compared to learning a transition model, because in the latter, model errors tend to accumulate during sequential resampling, making learning unstable. That said, the above theories are not mutually exclusive. Generative replay can be effectively used to simultaneously train the generative model and reactive controllers, as exemplified by DYNA or the world model approach (Ha and Schmidhuber, 2018). Furthermore, generative replay can directly support prospective functions of goal-directed controllers, such as the prediction of future paths. Therefore, our proposal is compatible with previous proposals that the hippocampus may operate synergistically with other brain structures, such as the ventral striatum, to form a goal-directed controller for spatial navigation (Pezzulo et al., 2014; van der Meer et al., 2012; Wikenheiser and Redish, 2015).

### Theoretical implications of the model

Casting the hippocampal formation as a hierarchical generative model may help reconcile its roles in multiple, apparently disconnected cognitive functions, including episodic and autobiographical memory (Eichenbaum, 2000; Tulving, 2002), prospective simulations and imagination (Buckner, 2010; Schacter et al., 2012) and cognitive map-based spatial navigation (O’keefe and Nadel, 1978; Tolman, 1948) which have often been studied in isolation from one another. Standard theories of system consolidation assume that episodic memories are transferred from the hippocampus to the cortex (McClelland et al., 1995; O’Reilly et al., 2014). However, a large body of evidence indicates that episodic memories may remain dependent on the hippocampus (Yonelinas et al., 2019). The theory proposed here suggests a specific way episodic memories may remain depend on the hippocampus: by being instantiated in the hippocampal generative model. Despite the generative model keeps changing with novel experiences, it is still capable to re-enact (with a varying level of fidelity) previous experiences. This idea is in keeping with the hypothesis of “constructive episodic simulation”, which suggests that episodic memory is an inherently constructive process and furthermore, that it provides details to construct simulations of the future - therefore explaining why episodic memories and prospective simulations share largely the same substrate and the same constructive processes (Schacter et al., 2007; Daniel L Schacter and Addis, 2007; Szpunar, 2010). Our model provides a mechanistic grounding to this hypothesis, by showing that generative replay supports both the re-enactment of episodic memories and prospective simulations / imagination, that is, the spontaneous generation of novel experiences that are structurally coherent with past experiences (as they stem from the same maps). Intriguingly, the theory of constructive episodic simulation suggests also that an imperfect memory that recombines previous experiences can be more useful to simulate future events compared to a memory that stores perfect records (Daniel L. Schacter and Addis, 2007). Our results support this idea by showing that generative replay confers systematic advantages in learning and generalization compared to the most widely used mechanism of experience replay, which epitomizes perfect memory storage.

Another important aspect of the model is that it organizes experiences into coherent spatiotemporal contexts. This makes it possible to discriminate between experiences that share similar features but occurred in different contexts, which is a key feature of episodic memory. Furthermore, we argue, the very same mechanism is crucial for cognitive map formation. As discussed above, the possibility to organize experiences within independent relational maps (at the highest level of our model hierarchy) confers significant advantages, especially during the continual learning of multiple tasks. In our model, the maps may automatically include structural priors, such as the fact that there are small gaps between adjacent locations and no preferred directionality, which implicitly embody characteristics of the physical space where an animal navigate - but more speculatively, may provide a relational scaffold to organize experiences in many more domains of cognition beyond spatial navigation (Behrens et al., 2018; Bellmund et al., 2018; Buzsáki and Moser, 2013; Constantinescu et al., 2016). Finally, and speculatively, the hippocampal generative model may contribute to autobiographical aspects of memory, as by continuously encoding and replaying episodes it may afford a continual sense of self that persists across time. This may also link to prospection, as a model provides not just a memory of “the past me” but also inference of “the future me” (Addis et al., 2011).

In sum, the computational framework of hierarchical generative modelling and generative replay reconciles several apparently disconnected theories of hippocampal function and provides a formal framework for their empirical testing.

### Future challenges

Though we illustrated our proposed model in the domain of spatial navigation, an interesting possibility that remains to be explored is that it may support the autonomous learning of coherent spatiotemporal contexts beyond navigation as well. Recent studies have shown that the hippocampus supports relational inference (Eichenbaum, 2000), temporal and rate coding of discrete sequences of non-spatial events such as (in rodents) odours (Terada et al., 2017) and button locations (Miller et al., 2017) and (in humans) of arbitrary items (Y. Liu et al., 2019); see also (Schuck and Niv, 2019; Shahbaba et al., 2019). More broadly, several findings imply a role for the hippocampal formation in cognitive mapping in several domains of cognition outside of spatial navigation (Behrens et al., 2018; Bellmund et al., 2018; Buzsáki and Moser, 2013). It remains to be established to what extent the hippocampus is a *general-purpose* system to form maps and sequences, and whether the generalization to non-spatial domains reuses internal codes initially developed for spatial navigation (Buzsáki and Moser, 2013). Theoretical arguments suggest that while the brain might use multiple generative models (G. Pezzulo et al., 2017), these may be tuned to different (natural) statistics of stimuli. At one extreme, generative models supporting visual processing may be tuned to scenes that change gradually (in the sense that successive frames tend to be similar to one another). The hippocampal generative model might lie close to the other extreme: it may be specialized to learn sequences of arbitrary (pattern-separated) items (Giovanni Pezzulo et al., 2017). Establishing the plausibility of these theoretical arguments is an objective for future research.

The model proposed here is simplified in many respects; for example, it does not fully include the rich inputs (e.g., sensory and spatial codes) that the hippocampus might receive from the entorhinal cortex. Future extensions of the model might consider these richer inputs and study in which ways they might shape the formation of spatial maps. Another possible extension of the current model would be including another prominent kind of internally generated hippocampal sequences: *theta sequences*. These are ordered sequences of place cell activation within each hippocampal theta cycle and occur during navigation (Foster and Wilson, 2007). Theta sequences have been implied in both memory encoding and the prediction of future spatial positions along the animal’s current spatial trajectory (Johnson and Redish, 2007; Lisman and Redish, 2009; Redish, 2016). It is possible that theta sequences are yet another manifestation of the same hippocampal generative model that supports replay, which arises spontaneously when an animal is task-engaged (Giovanni Pezzulo et al., 2017). If this were the case, it would be possible to implement theta sequences in our model, by rapidly interleaving the (bottom-up) recognition of the current sequence and the (top-down) generation of predicted future locations, within the same theta cycle. This implementation would follow the proposal that early and late phases of each theta cycle support the inference of the current state and the prediction of future states, respectively (Sanders et al., 2015). In other words, the first part of a theta cycle would play the role of a “filter” (requiring bottom-up inputs) to represent the previous and current location, whereas the second half of the cycle would play the role of a “predictor” (requiring top-down inputs) to represent future locations. A speculative possibility is that the theta rhythm itself acts as a gain modulator for bottom-up streams (Shimazaki, 2018), by increasing their precision during the first half of each theta cycle and then decreasing it during the second half – thus only allowing top-down predictions to occur in this second half. These or alternative implementations of theta sequences remain to be tested in future studies.

A potential challenge to this model is that hippocampal reactivations do not always show the diffusive dynamics reported in (Stella et al., 2019) and analysed in this article. Various studies have reported that trajectory events tend to “jump” in the awake behaving state (Krause and Drugowitsch, 2022; Pfeiffer and Foster, 2015, 2013). In our model, the reactivation dynamics (diffusive or otherwise) mainly depend on a displacement coefficient (*c*) parameter that controls sequence continuity. Following (McNamee et al., 2021), it is possible to hypothesize that a modulation of grid cell activity may alter the statistical structure of hippocampal sequence generation - which, in our model, is achieved by changing the displacement coefficient parameter. This hypothesis remains to be tested in future studies.

## Methods

### Hierarchical spatiotemporal representation

Our theory proposes that the hippocampus implements a hierarchical generative model that organizes the *observations* it receives over time into three sets of latent codes, for *items*, *sequences* and *maps*, respectively. At the highest hierarchical level, the model learns a set of maps representing characteristic environmental (e.g., maze) properties. The model implicitly assumes that, at each time (or short time interval), its observations derive from one single map that must be inferred. A key characteristic of the generative model is that it inherently implements *generative replay*: it can stochastically generate sequences of items from the inferred map; and use the stochastically generated sequences to train the model itself (or support other cognitive task, such as trajectory prediction and planning).

Before formally describing the model, we need to introduce some notation. Let us assume that the environment is represented as a vector space 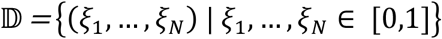 of dimension *N*, which in our simulations of spatial navigation correspond to a domain of *N* spatial locations. An *observation* is defined as a one-hot vector: an event of “magnitude” *ξ_i_* occurring at a location *i* = 1, …, *N*. Using the standard basis **e**_1_ = (1, 0, …, 0), **e**_2_ = (0, 1, …, 0), …, **e**_*i*_ = (0, …, 1, … 0), **e**_*N*_ = (0, 0, …, 1), observations are represented at the lowest level of the hierarchical model as ***ξ**_i_* = *ξ_i_***e**_*i*_. The default magnitude *ξ_i_* of a noiseless observation at position *i* is set to 1.

Moving upward, the next hierarchical level infers the hidden code **x** of the observations in 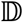, called *items*, such that

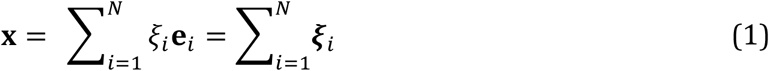

Moving upward again, the next hierarchical level uses the same encodings to define a hidden code **y** for a *sequence* of items *S* ≡ ***ξ***(1), …, ***ξ***(*T*) representing a series of observations, or a trajectory through 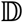:

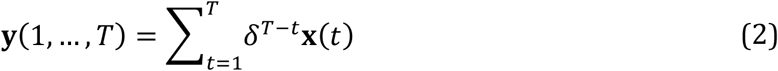

where **0** < *δ* < 1 (*δ* = 0.7 in our simulations) and *δ^T–t^* is an exponential decay over time. Equation (2) consists of a graded representation (or trace), where the most recent observation is encoded with the highest activation rate. Note that this coding scheme closely resembles a “bag-of-words” encoding of multiple words (observations) in a document (sequence). However, while the bag-of-word representation encodes the frequency (distribution) of elements without any order (*δ* = 1), our method assumes few repetitions such that the graded values implicitly encode item order.

Finally, at the top level, the model assumes that sequences of items **y** can be grouped together, as if they were drawn from a distribution that partitions the sequences into clusters. Each cluster *k* corresponds to (and is characterized by) a different feature map described by a vector of parameters 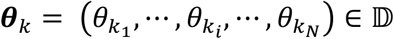 encoding the frequency of event occupancy of the 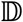 locations under map *k*. Thus, the model assumes that the representation **y** of a sequence of observations is drawn from cluster *k* according to a probability *p*(**y**|*θ_k_*) and that the prior probability of this cluster is *p*(*k*). These assumptions are embodied in the following generative *mixture model*:

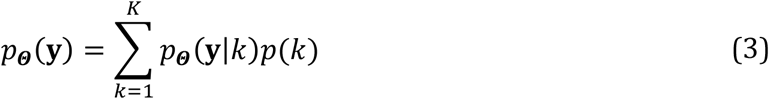

where 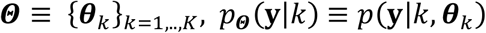, and *K* is the number of clusters. Note that while here we set *K* as known and fixed, in the section “Nonparametric Method” we discuss a non-parametric extension of the model, which automatically infers the number of clusters given its observations.

The definition of the model in Equation (3) is well-posed once we specify the set of mixing probabilities *p*(*k*) and the conditional distributions *p_**Θ**_*(**y**|*k*). In the proposed model, we assume that ***z*** ≡ (*p*(*k* = 1), …, *p*(*k* = *K*))~*Cat*(***π***) and the parameters ***θ**_k_*~*Cat*(***ρ**_k_*) with ***ρ**_k_*~*Dir*(1, …,*N*). Therefore, both the mixing probabilities and the structural parameters of the maps follow a categorical distribution – the latter having hyperparameters drawn from a Dirichlet distribution over 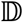.

In turn, we define the likelihood *p_**Θ**_*(**y**|*k*) that the *k*^th^ map can generate the representation **y** of sequence of observations *S* as a deterministic function of **y** and the probabilistic parameters ***θ**_k_* of that map:

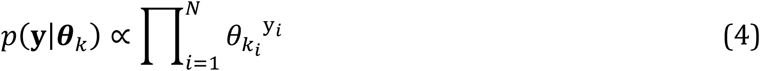

where y_*i*_ = **y** ∘ **e**_*i*_, with the symbol “**∘**” denoting the Hadamard (element-wise) product. By taking the logarithm, Equation (4) becomes the scalar product between **y** and ln ***θ**_k_*:

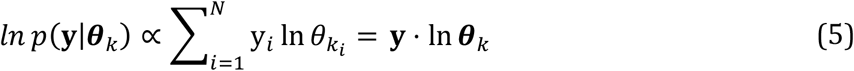

Equations (4) and (5) effectively measure the compatibility of the sequence of events under the structure of a given map, on a linear and a logarithmic scale, respectively. Indeed, equation (5) assigns higher probabilities to those maps whose spatiotemporal structure ***θ**_k_* more closely matches the allocation and temporal order of observations encoded by **y**. Note that in equations 4 and 5, the likelihood of sequences does not explicitly take into account the temporal order or the adjacency of items, but only the occupancy frequency in a maze. However, as explained above, temporal information is implicitly encoded in the sequence representation, given that the activation of past items decays with time.

A final thing to notice is that Equation (5) affords a plausible neuronal implementation in the mechanism of weighted transmission of action potentials between synapses encoding log-probabilities of the spatiotemporal structure and presynaptic neuronal activity encoding event representations.

### Hierarchical inference

To infer the posterior of the latent states upon the observation of a new event, we adopt an approximate inferential approach that considers the hierarchical and temporal dependencies of the model (Figure 1a). Please see the “**`inference`**” procedure in the Supplementary Materials for more details.

First, we estimate the posterior of the mixing probability distribution ***z***(*t*) ≡ *p*(*k*|**y**(*t*). This is done differently at the beginning of an episode (when *t* = 0) and afterwards (when *t* > 0). At the beginning of an episode, the mixing prior ***z***(0) is drawn from a categorical distribution ***z***(0)~*Cat*(***π***) with hyperparameters ***π*** that follow a Dirichlet distribution, ***π**~Dir*(***α***). After the calculation of the posteriors, we recursively adjust the hyperparameters ***π*** based on the accumulated evidence, thus allowing agents to improve the a-priori information available for the next trials; see the section on “Parameter learning” for details.

In contrast, when *t* > 0 the posterior of ***z***(*t*) is calculated as follows:

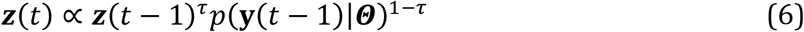

Here, *τ* is an exponential time constant, whose value is fixed to (the empirically determined value) *τ* = 0.9 and ***Θ*** is the full set of maps ***θ**_k_*. For the sake of simplicity, we recall Equation (5) and use a log-scale to change Equation (6) to the linear expression:

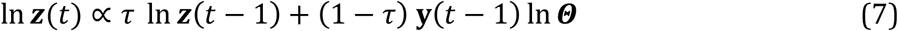

where the matrix multiplication **y**(*t* – 1) ln ***Θ*** sums over the spatial domain.

Equation (6) determines the posterior mixing probability **z**_*k*_(*t*) of each cluster *k* at the current time step by dynamically integrating its prior state **z**_*k*_(*t* – 1) and the likelihood of the current sequence, which considers the compatibility between **y**(*t* – 1) and the cluster. We adopt a maximum a posteriori (MAP) approach to select (and adapt) which of the *K* maps maximizes the posterior; namely 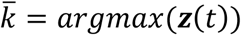. By doing so, latent state inference guides learning to gradually form distinct maps for different environments.

At the end of the inferential process at each time step, the parameters 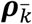 that define the selected map 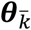 are updated; see the section on “Parameter learning” for details.

Note that Equation (6) can be interpreted as Bayesian filtering, by considering that it operates in two sequential steps: generative prediction and Bayesian update. We could interpret Equations 6 and 7 as the probabilistic prediction of a set of *K* hidden states *k* = 1 … *K* conditioned on the sequence code **y**(*t* – 1). However, unlike standard Bayesian filtering, the latter is another latent state, not an observation. This process dynamically reduces uncertainty in map distributions (and the entropy of ***z***(*t*)), hence the inference provides increasingly more precise estimates of the mixing probabilities and map inference. This first inferential step has a plausible biological implementation, in terms of a local competition circuit that recursively accumulates evidence for each map, achieved by adding synapse-weighted bottom-up signals encoding current observations at each time step.

Second, we estimate the posterior item codes **x**(*t*). (Note that the hierarchal structure shown in Figure 1A might suggest that the posterior sequence **y** should be estimated before the posterior item codes **x**(*t*). However, we do the opposite to simplify the generation of fictive information during replay, see below). The dynamics equation for **x** considers three elements: (i) the map 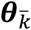 inferred at the highest hierarchical level, (ii) a simple physical transition model specifying that the transitions are sampled from a normal distribution centred on past observation ***ξ***(*t* – 1) or in other words that only adjacent locations (in 2D space) can be visited next and (iii) the prior sequence code calculated at the previous time step **y**(*t* – 1), which is subtracted, thus implementing a form of “inhibition of return”. Thus, we define the posterior of **x**(*t*) as follows:

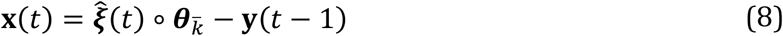

where 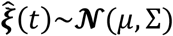 is a noisy prediction about the observation with a normal distribution centered on *μ* = ***ξ***(*t* – 1) + *c* * ***v***(*t* – 1) and with symmetric covariance |Σ| = ***v***(*t* – 1) modulated by the current velocity (i.e., the velocity at the tip) of the sequence **y**, ***v***(t) = ***ξ***(*t*) – ***ξ***(*t* – 1) and by an arbitrary chosen coefficient *c* = 0.4 promoting sequence continuity.

After updating the item code **x**(*t*), the model generates a prediction for a new observation through the maximization of the inferred item, i.e.:

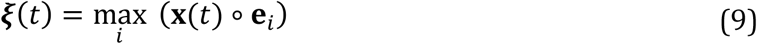

This mechanism is key for generative replay and the generation of fictive observations over time.

Third, we estimate the posterior sequence code **y**(*t*), using the following dynamics equation, which considers the hierarchical structure and causal temporal dynamics of the model:

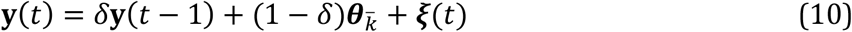

where *δ* is the decay coefficient already introduced in Equation (2). In practice, Equation (10) gradually creates a spatiotemporal representation of observed sequences, by adding the item code ***ξ***(*t*) corresponding to the current observation to the previous sequence code **y**(*t* – 1), while also considering the top-down prediction provided by the inferred map 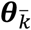. Note that the item code ***ξ***(*t*) is inferred during navigation, but fictive during replays.

### Parameter estimation (learning)

In parallel to the aforementioned state-inference process, the model continuously infers its (hyper)parameters. In particular, every time the latent-state inference selects the *k^th^* hidden map via Bayesian filtering, it updates the model hyperparameters 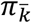 and 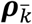. This enables long-term environmental information (e.g., maze properties) to be maintained across time. Please see the “**`learning`**” procedure in the Supplementary Materials for more details.

To improve hyperparameter learning, we adopt the following methods: (i) we use a (small) decay constant *γ* to account for the volatility of information coded in the mixing priors (in our simulation, this is set to 1%), (ii) we scale the increments of the Dirichlet counters with the probability 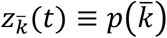 to account for the confidence in choosing 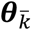 as the best map. This update is done after the hierarchical inference step.

To calculate the posterior of the hyperparameters of the 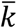 mixing probability 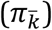, we use the following update:

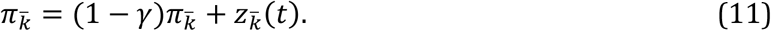

Note that even if the value of 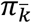 is updated at every time step, it is only used at the beginning of each trial (*t* = 1), as a parameter of the categorical distribution from which the prior ***z***(0) is sampled. Therefore, equation (11) embodies a-priori information (from previous trials) about the probability of each cluster in the parameters ***π***.

To infer the Dirichlet hyperparameters of the selected map 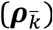, we use the following:

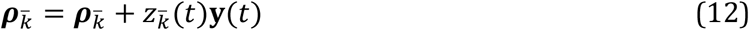

which accounts for both the temporal information in the sequence code **y** and belief 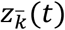 of the choice of the current map. This update is done only when t > 1, to estimate the most likely map associated with the sequence of observations observed so far.

Finally, we generate the *prioritized map* 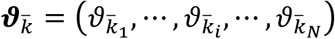 used by the prioritized generative replay agent (see the section *“Training Procedure”*) by considering the surprisal of observations ***ξ*** during the inference-state process at every time step *t* and following the following recursive expression:

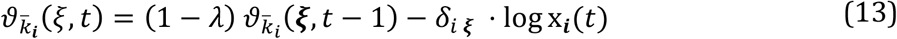

In the equation, ***δ**_i ξ_* denotes the Kronecker delta function, which is 1 if *i* and *ξ* are equal and 0 otherwise; *λ* is a small decay factor with value in the unit interval; and x_*i*_ is the posterior probability given by Equation (8). It is worth noting that the map defined in Equation (13) takes larger values on unlikely observations *ξ* following the notion of Shannon surprise.

### Nonparametric method

The simulations described so far used a fixed number of mixture clusters *K*. Here we describe an extension of the previous model that automatically infers *K*. For this, we use an adaptation of the Chinese Restaurant Process (CRP) prior over clustering of observations that adapts model complexity to data by creating new clusters as the model observes novel data (Gershman and Blei, 2012; Stoianov et al., 2015). Note that the CRP prior is applied only during navigation, on true data, and not during replay. This is done to prevent the generation of an excessive number of clusters.

The CRP metaphor describes how a restaurant with an infinite number of tables (clusters) is gradually filled in with a potentially infinite number of customers (observations). The first customer sits at the first table, i.e., it is assigned to the first cluster. All the other new customers that enter the restaurant can either sit at an occupied table (i.e., be assigned to existing clusters) with a probability corresponding to table occupancy *n_i_*/(*n* + *α*) or sit at a new table (i.e., expand the model with a new cluster) with probability *α*/(*n* + *α*), where *α* is a concentration parameter.

Notably, our customers are multi-dimensional real-value representations **y** and each representation is different from any of the previously observed ones. To determine whether a representation **y** should be considered as novel, we extended the classical CRP prior method to also exploit the knowledge stored in the learned cluster parameters: the maps ***θ**_k_*. A representation **y** is considered as novel when the greatest likelihood *p*(**y**|***θ**_k_*) (see Formula 4) among all clusters currently in use is smaller than a threshold, *p_thr_* = 0.01. Table occupancy corresponds to the parameters *π_k_* of the mixing probabilities and the concentration parameter is *α* = 1.

### Rank-maps

Rank-order maps are used in (K. Liu et al., 2019) to characterize the similarity between pre-run (preplay) and post-run (replay), and between post-run and run (navigation experience). We calculate the rank map of each navigation or replay episode by accumulating the spatial trajectories 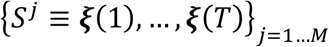 represented in terms of normalized localized ranks during the episode. Thus, for a given episode, the rank map **R** is defined as follows:

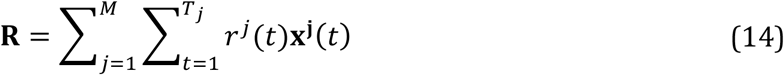

where 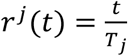 is the normalized rank at time **t** of sequence *j, T_j_* is the length of sequence *j*, and **x^j^**(*t*) is the location at time *t* of sequence *j* (defined in Equation 1). For replay episodes the predicted observation is used instead of the true observation (which is not available). Finally, Spearman’s rank order correlation is used to calculate the similarity between the (vectorized) ranks maps. Examples of rank maps and rank correlations are shown in Fig. 6A and statistics of rank-correlations across conditions is shown in Fig. 6B.

### Dynamics of Diffusion

To characterize the trajectory dynamics during navigation and replay, for each learner from each training condition during either navigation or replay, we collect the temporal (Δ*t*) and Euclidean (L2) spatial distance Δ*x* between each pair of points *ξ*_*t*_1__ and *ξ*_*t*_2__ (chunks) from each trajectory *S* ≡ ***ξ***(1) … ***ξ***(*T*). We then fit a log-log regression model of the spatial distance as a function of temporal distance (i.e., duration): ln(Δ*x*) ≈ *α* ln (Δ*t*). Note that the estimated regression coefficient α directly corresponds to the exponential coefficient of the power law: Δ*x*=Δ*t^**α**^*.

The estimation is applied on all trajectory chunks, up to a limit of 20 time steps long (this limit it selected given that the map size is 15×15 units; longer sequences would necessarily involve some systematic curvature) The results of the analyses are shown in Figure 4A.

Finally, we collect the spatial statistics of the distribution of diffusion relative to the initial position of each chunk. Some examples are shown in Figure 4B.

## Acknowledgements

This research received funding from the European Union’s Horizon 2020 Framework Programme for Research and Innovation under the Specific Grant Agreement No 945539 (Human Brain Project SGA3) to GP, the European Research Council under the Grant Agreement No. 820213 (ThinkAhead) to GP, the European Union’s Horizon H2020-EIC-FETPROACT-2019 Programme for Research and Innovation under Grant Agreement 951910 to IS and from the Italian Ministry for Research MIUR under Grant Agreement PRIN 2017KZNZLN to I.S. We gratefully acknowledge Jeremy Gordon for useful comments on this manuscript.

## Supplementary Materials

### Control simulations

We ran two sets of control simulations aiming to replicate the main simulation and investigate the robustness of the model and the effect of two parameters (number of replays, and item sampling error). Each simulation included n=32 replicas per condition.

First, we investigated the role of the number of replays by using twice, four times, half, and a quarter the number of navigation trials in a single episode (for which there were n=20). As in the main simulation, model evaluation was based on the proportion of incorrectly recognized mazes during the two phases of learning: navigation and retention. The results shown in Table S1 confirmed the overall advantage of generative replay methods, which (as expected) gradually reduces with fewer replays. However, using more replays did not consistently improve the performance, and in some cases slightly worsened it. This result may be due to the fact that an excessive strengthening of memory traces might decrease choice flexibility.

Second, we investigated the role of the displacement coefficient *c* that controls sequence continuity in formula (8) by using either half or double its value (*c* = 0.4 in the main simulation). Greater displacement was expected to result in less consistent sequences and therefore decreased map precision and reduced performance. The results shown in Table 1 confirmed that intuition. Doubling displacement (i.e., *c* = 0.8) decreased the advantage of replay and halving it (i.e., s=0.2) did not show consistent performance improvement.

Figure S5 shows the maps of sample learners in several conditions. These maps confirm the model robustness and suggest that lowering the number of replays (to n=5) and increasing the displacement coefficient (c=0.8) results in less consistent maps and greater categorization error, which reduces accuracy in item- and map-inference. For example, the sample learners in the condition of large displacement coefficient (s=0.8, bottom row) have the same maps for maze 3 and 4, which is an example of categorization error: the same cluster has become preferred for two very similar mazes.

### Supplementary Video

We have provided a supplementary video exemplifying the functioning of the hierarchical generative model discussed in this article. The video shows the model inputs (bottom panel) and the activity of the latent units at each hierarchical level, during the first three and the last three trials of each of the five learning blocks. For illustrative purposes, the video shows the dynamics of generative replays after each trial (labelled as “awake replays”), but these are not used for learning. Furthermore, the video shows the dynamics of generative replays after each block (labeled as “replays during sleep”), which are used for learning. The video illustrates that the hierarchical generative model learns accurate maps very rapidly, which allows accurate inference of which maze the agent is currently in. Furthermore, the generative replay mechanism generates both sequences that were actually experienced and novel sequences (sometimes including implausible sequences, as expected).

**Supplementary Table S1.** Control simulations. Recognition error measured during navigation (learning) and retention test. The gray rows are two replicas of the main learning condition (20 replays, s=0.4). There are n=32 replicas per condition. Replays denote the number of replays and c denotes a displacement coefficient parameter that controls sequence continuity. See the main text for explanation.

| LEARNING |  |  |  |  |  |  |  |  |  |
| --- | --- | --- | --- | --- | --- | --- | --- | --- | --- |
| replays |  | Baseline |  | Exp.Replay |  | Gen.Replay |  | Prior.Gen.Replay |  |
|  | 5 | 0.16 | (se 0.02) | 0.14 | (se 0.02) | 0.11 | (se 0.02) | 0.12 | (se 0.01) |
|  | 10 | 0.14 | (se 0.02) | 0.12 | (se 0.02) | 0.11 | (se 0.02) | 0.11 | (se 0.02) |
|  | 20 | 0.16 | (se 0.02) | 0.10 | (se 0.02) | 0.07 | (se 0.01) | 0.05 | (se 0.02) |
|  | 40 | 0.17 | (se 0.02) | 0.12 | (se 0.02) | 0.08 | (se 0.01) | 0.03 | (se 0.01) |
|  | 80 | 0.16 | (se 0.02) | 0.12 | (se 0.02) | 0.08 | (se 0.02) | 0.05 | (se 0.02) |
| c | 0,2 | 0.14 | (se 0.02) | 0.13 | (se 0.02) | 0.05 | (se 0.02) | 0.07 | (se 0.02) |
|  | 0,4 | 0.16 | (se 0.02) | 0.10 | (se 0.02) | 0.07 | (se 0.01) | 0.05 | (se 0.02) |
|  | 0,8 | 0.16 | (se 0.03) | 0.12 | (se 0.02) | 0.08 | (se 0.02) | 0.15 | (se 0.03) |

| RETENTION |  |  |  |  |  |  |  |  |  |
| --- | --- | --- | --- | --- | --- | --- | --- | --- | --- |
| replays |  | Baseline |  | Exp.Replay |  | Gen.Replay |  | Prior.Gen.Replay |  |
|  | 5 | 0.21 | (se 0.02) | 0.20 | (se 0.02) | 0.17 | (se 0.02) | 0.18 | (se 0.01) |
|  | 10 | 0.19 | (se 0.02) | 0.19 | (se 0.02) | 0.17 | (se 0.02) | 0.17 | (se 0.02) |
|  | 20 | 0.22 | (se 0.02) | 0.18 | (se 0.02) | 0.14 | (se 0.01) | 0.12 | (se 0.01) |
|  | 40 | 0.22 | (se 0.01) | 0.21 | (se 0.01) | 0.15 | (se 0.01) | 0.11 | (se 0.01) |
|  | 80 | 0.21 | (se 0.02) | 0.22 | (se 0.02) | 0.17 | (se 0.02) | 0.14 | (se 0.02) |
| c | 0,2 | 0.20 | (se 0.02) | 0.21 | (se 0.02) | 0.13 | (se 0.02) | 0.13 | (se 0.02) |
|  | 0,4 | 0.22 | (se 0.02) | 0.18 | (se 0.02) | 0.14 | (se 0.01) | 0.12 | (se 0.01) |
|  | 0,8 | 0.22 | (se 0.02) | 0.20 | (se 0.02) | 0.14 | (se 0.02) | 0.20 | (se 0.02) |

### Supplementary Figures

**Figure S1.**
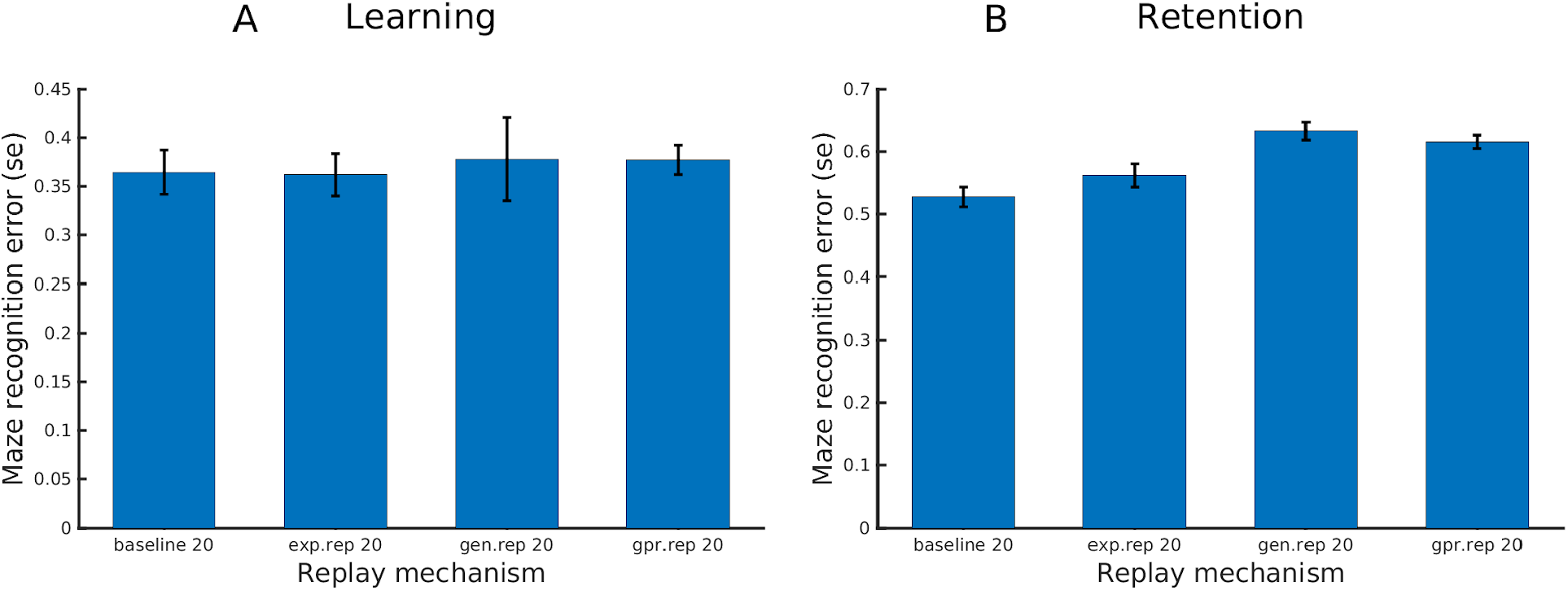
Performance of four agents (baseline, experience replay, generative replay, prioritized generative replay) that use a reduced model, which lacks the intermediate (sequence) layer. (A) Learning test. (B) Retention test. For comparison, the performance of the same four agents that use the full hierarchical model is shown in Figure 1C-D. See the main text for explanation. We found that in the Learning task, all the four agents perform significantly better when they use the full hierarchical architecture compared to the reduced model (baseline: t=4.612, sd=0.097, df=30, p=0.000; experience replay: t=6.386, sd=0.107, df=30, p=0.000; generative replay: t=5.958, sd=0.138, df=30, p=0.000; prioritized generative replay: t=11.960, sd=0.073, df=30, p=0.000). The same pattern emerges in the Retention task (baseline: t=10.102, sd=0.075, df=30, p=0.000; experience replay: t=11.894, sd=0.086, df=30, p=0.000; generative replay: t=17.877, sd=0.076, df=30, p=0.000; prioritized generative replay: t=21.026, sd=0.064, df=30, p=0.000). See the main text for explanation.

**Figure S2.**
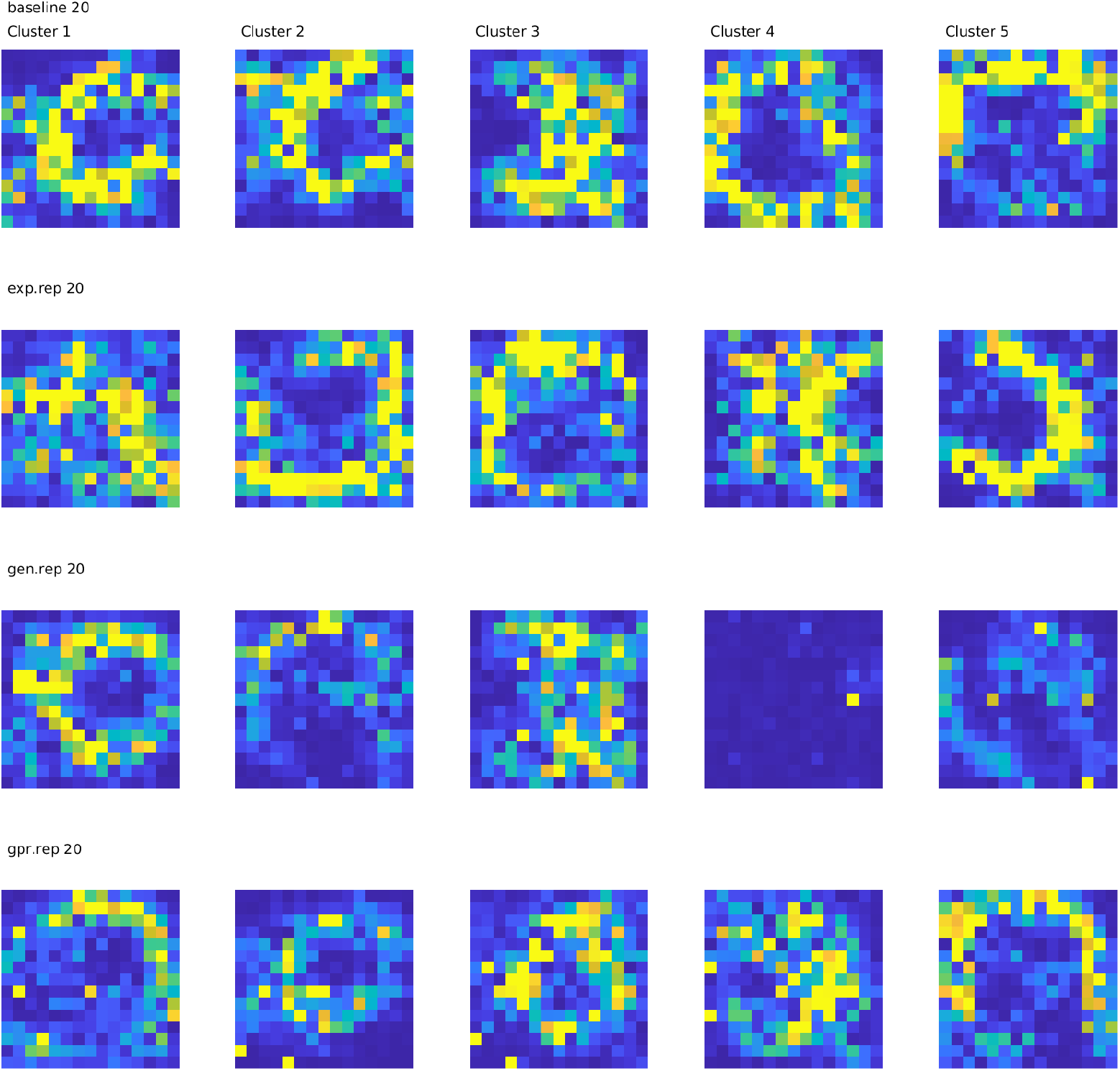
Maps learned by four sample agents that use a reduced model, which lacks the intermediate (sequence) layer. See the main text for explanation.

**Figure S3.**
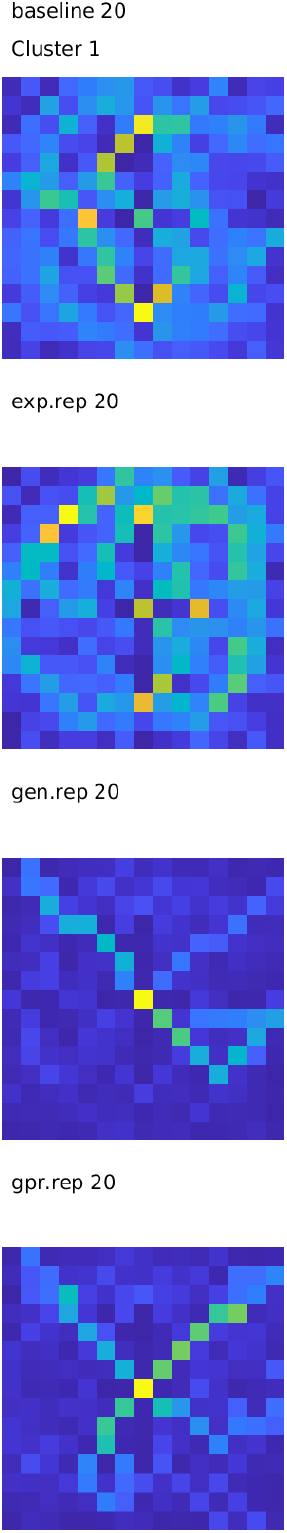
Maps learned by four sample agents that use a reduced model, which only includes 1 cluster at the highest (map) layer. See the main text for explanation.

**Figure S4.**
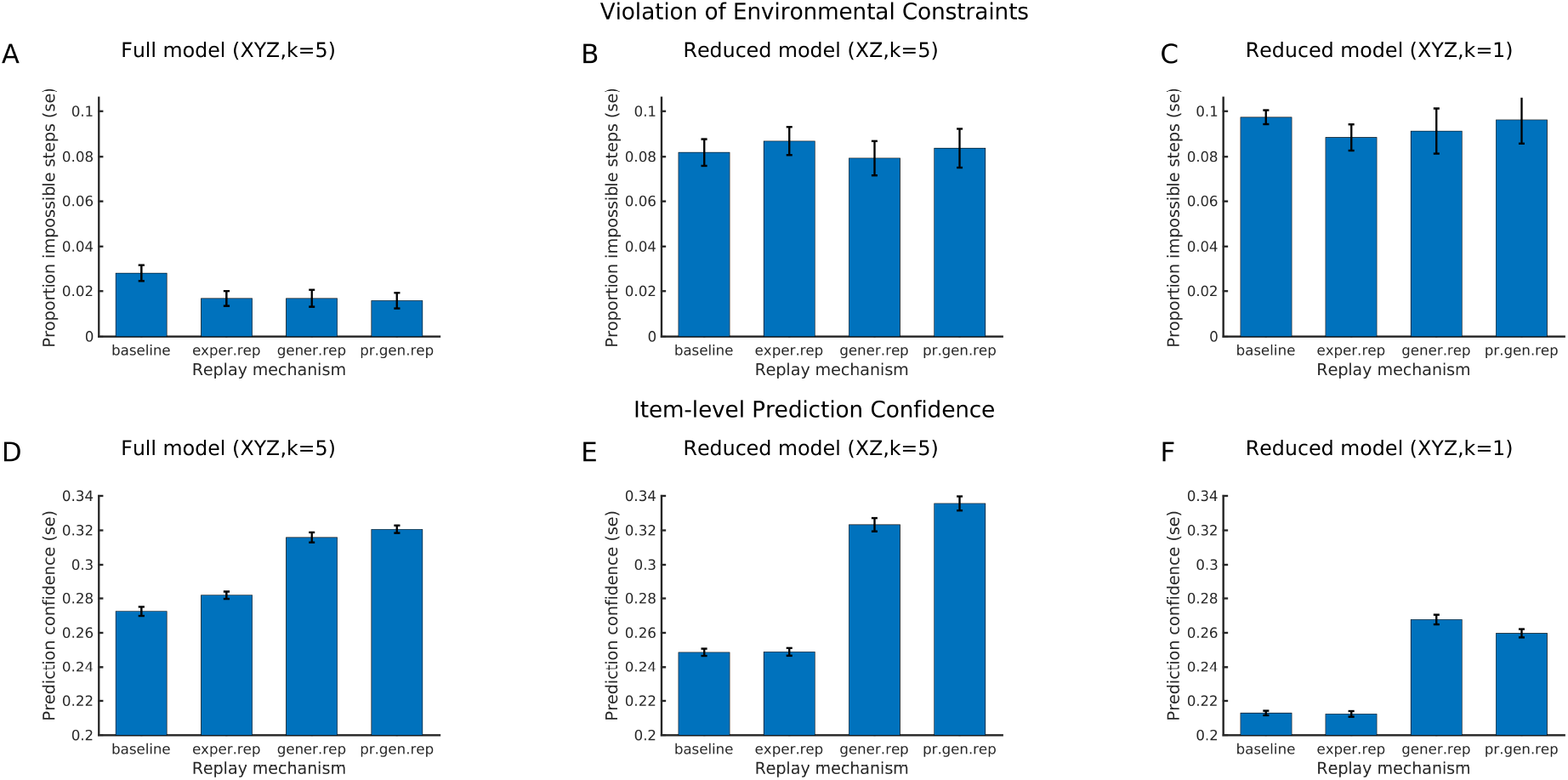
Comparison of the hierarchical generative model with reduced versions of the model that lack hierarchical depth. (A-C) Results of the test of violation of environmental constraints, for the full model, the reduced model that lacks the sequence layer and the reduced model that only includes 1 cluster at the highest (map) layer, respectively. (D-F) Results of the test of item-level prediction confidence. See the main text for explanation.

**Figure S5.**
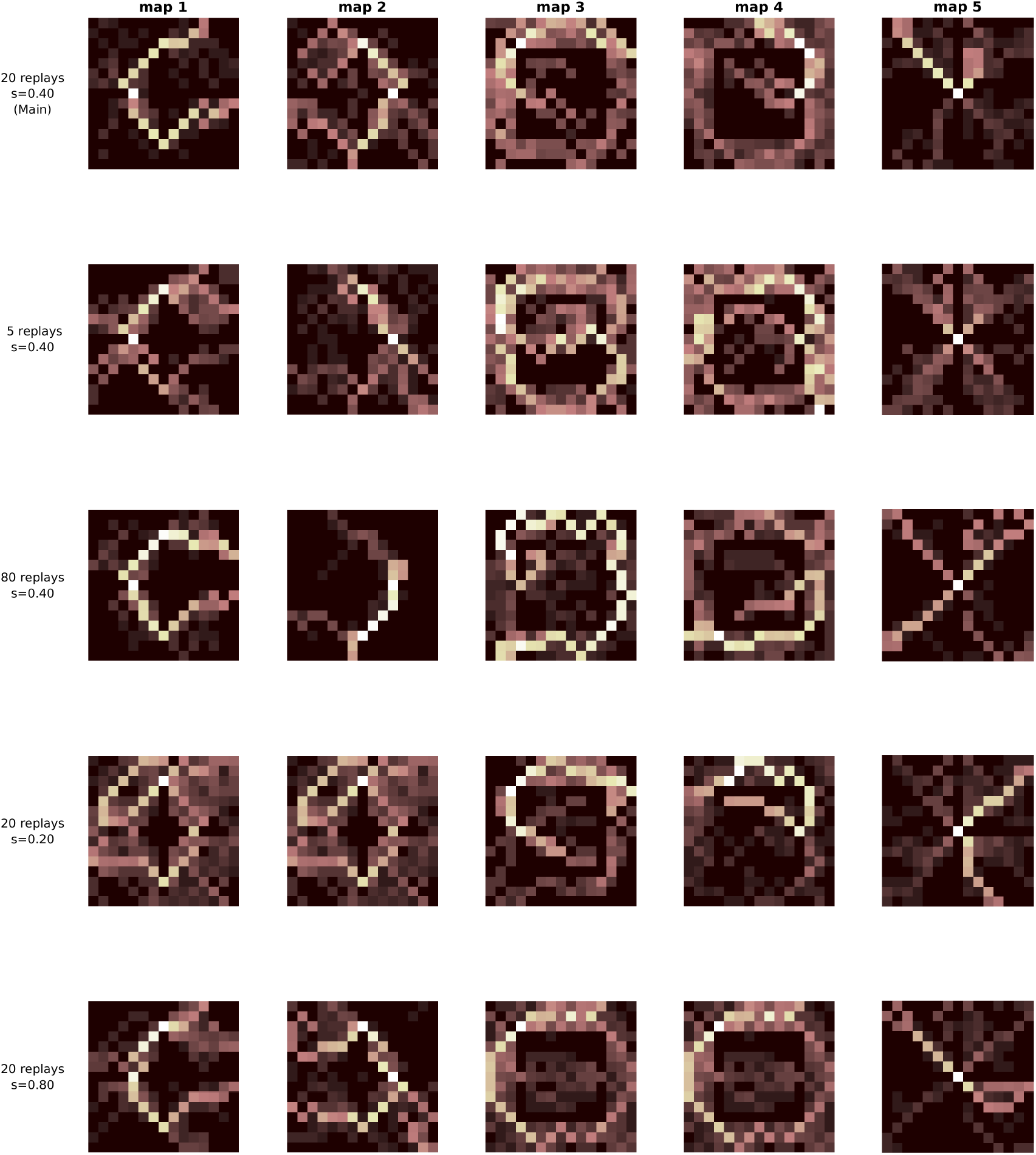
Maps learned by five sample learners that perform a different number of replays and use different displacement coefficients, as specified in Table S1. See the main text for explanation.

**Figure S6.**
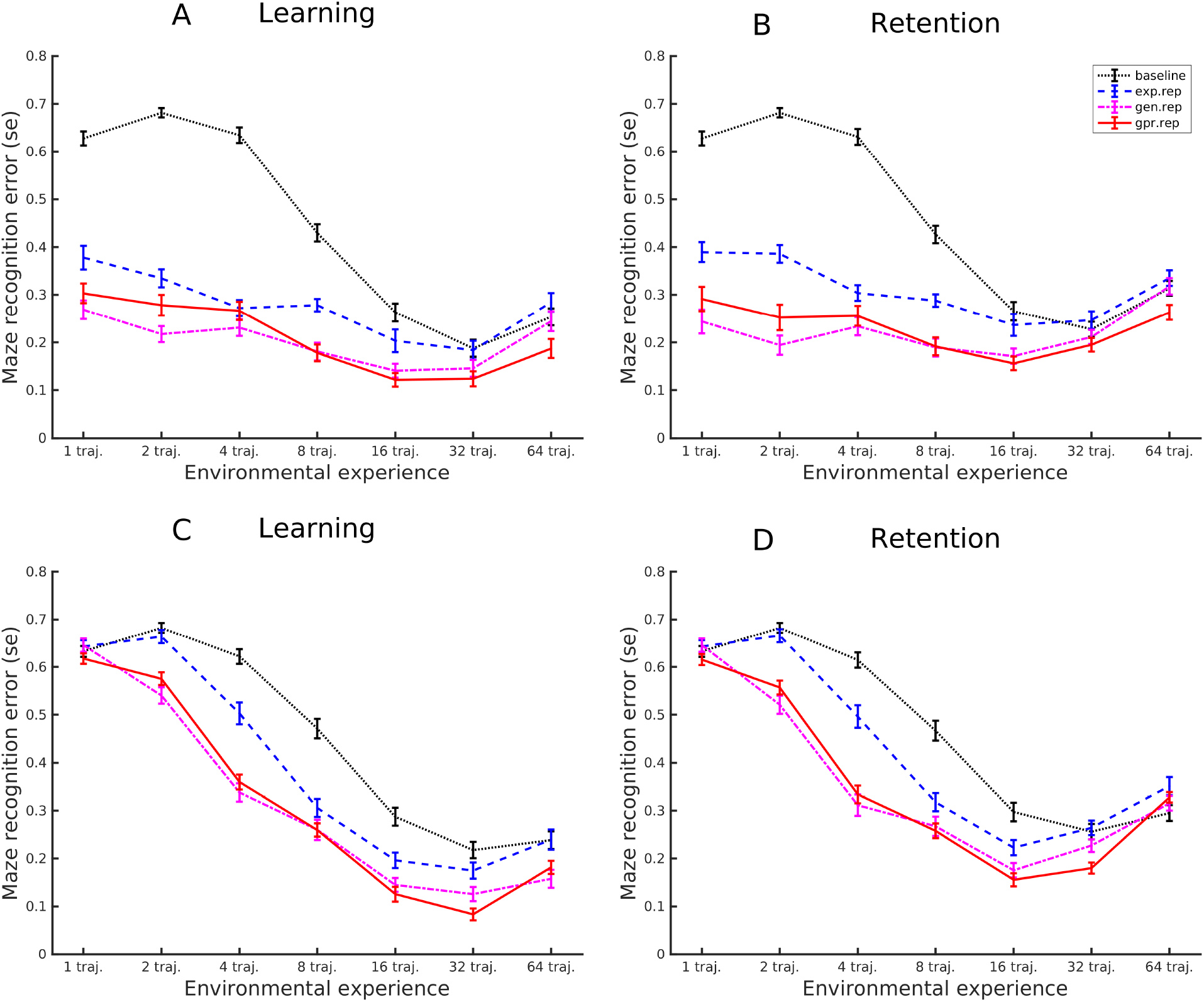
Control simulation, in which we varied the number of trajectories used as inputs during navigation (1, 2, 4, 8, 16, 32 or 64) and the number of replays executed afterwards. The plots show the performance of the baseline, experience replay, generative replay and prioritized generative replay agents during Learning and Retention tests, when the three replay agents replayed 20 trajectories after each block (A-B) or replayed the same number of trajectories as the number of inputs they received (e.g., when they received 1 input, they replayed 1 trajectory). See the main text for explanation.

**Figure S7.**
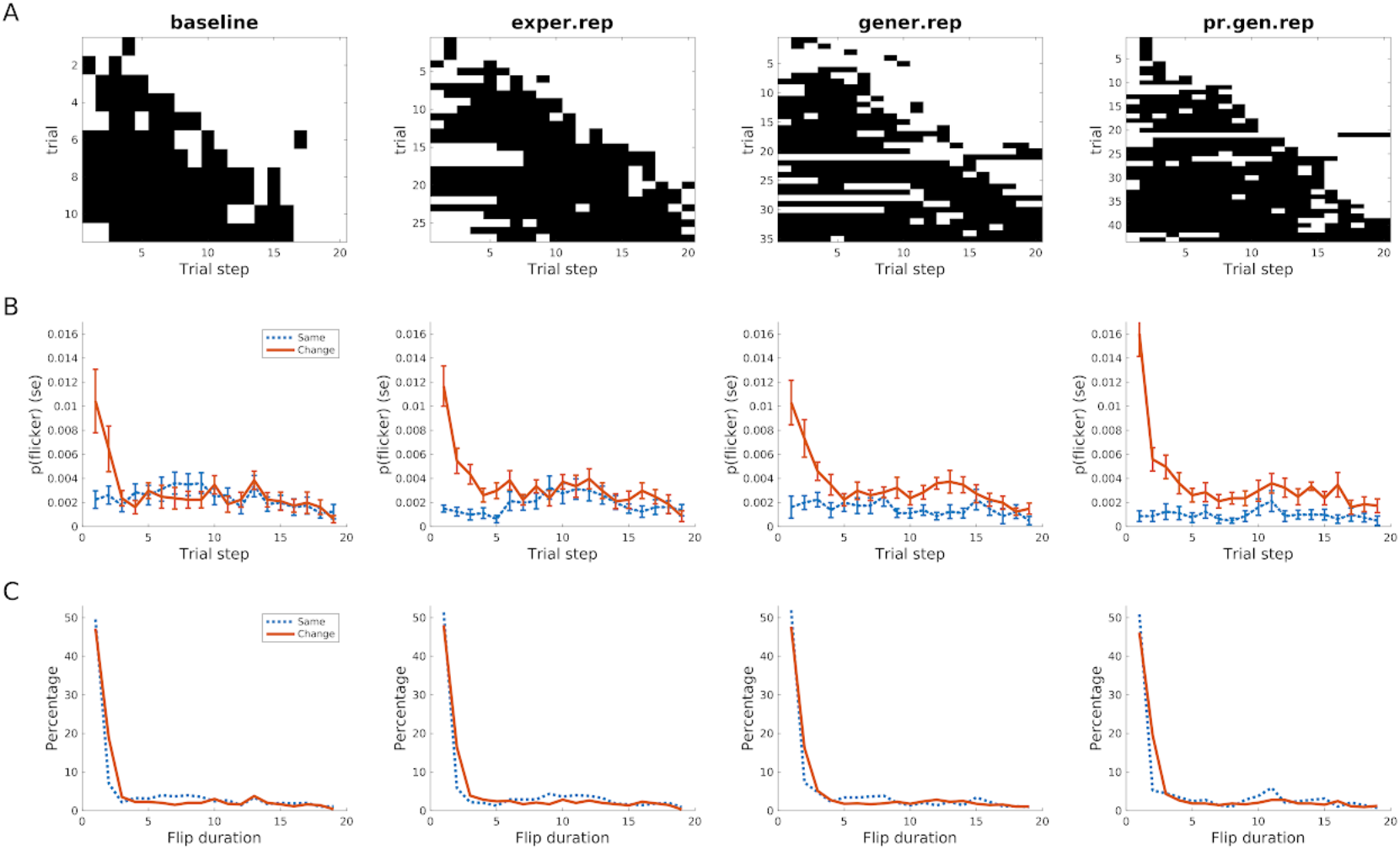
Simulation of flickering. (A) Example trials in which flickering occurs in the four agents. Flickering here corresponds to a sequence formed by a white square (i.e., correct inference) and then one or more black squares (i.e., incorrect inference). (B) Probability of flickering in the “same” and “change” conditions. (C) Duration of the flickering in the “same” and “change” conditions. See the main text for explanation.

### Pseudocode of the inference and learning procedures described in the Methods section

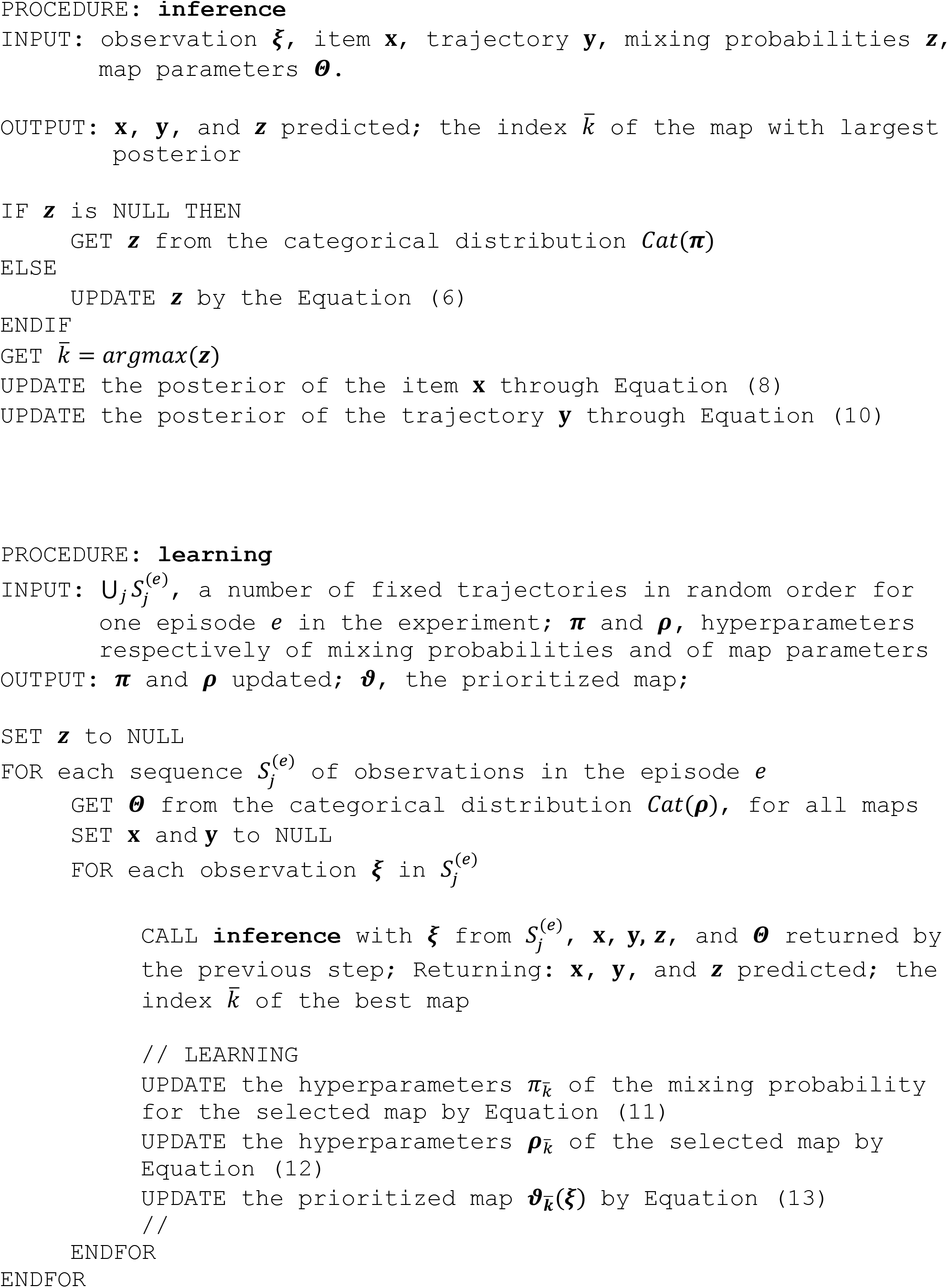

### Pseudocode of the four agents described in the main text

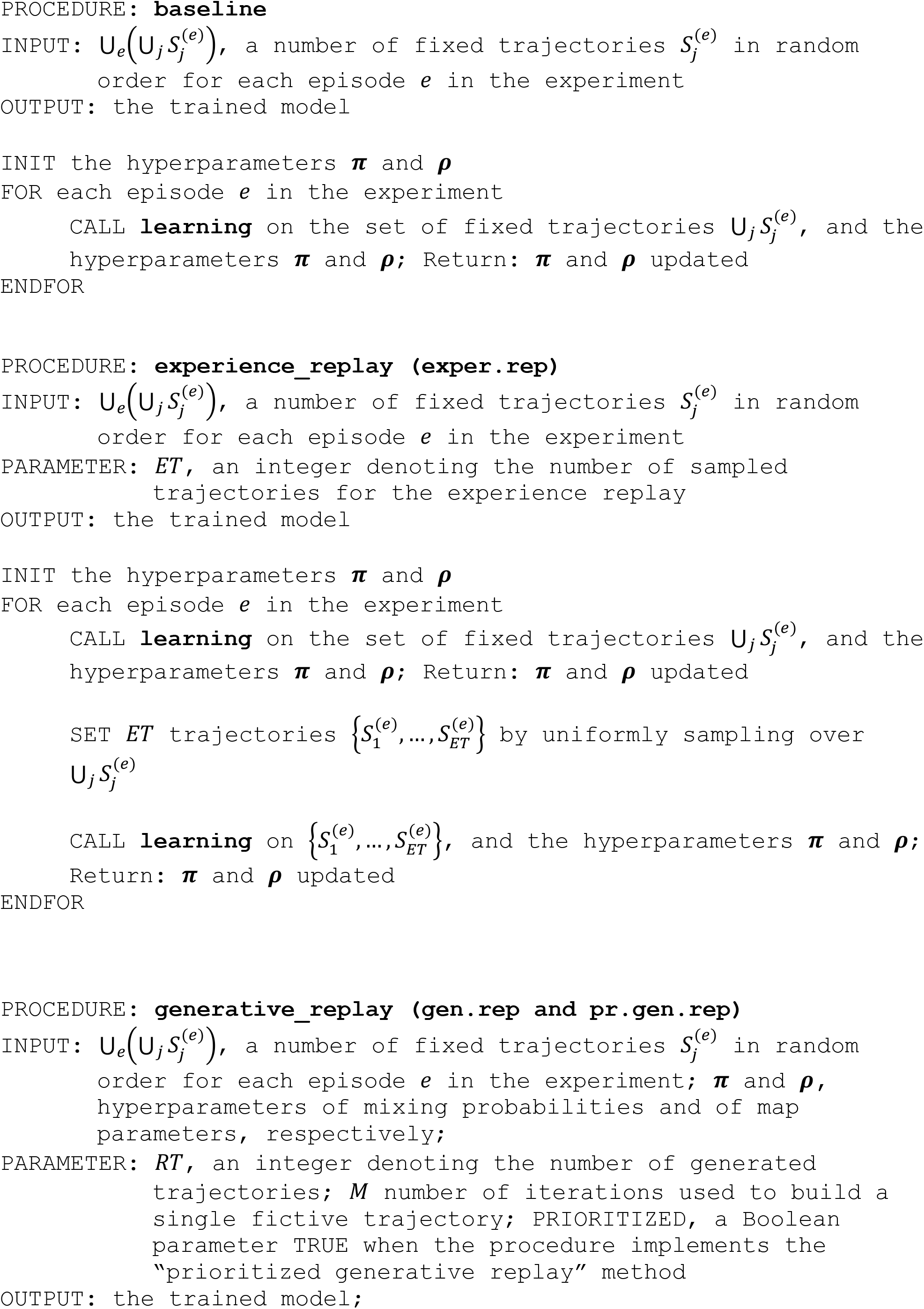
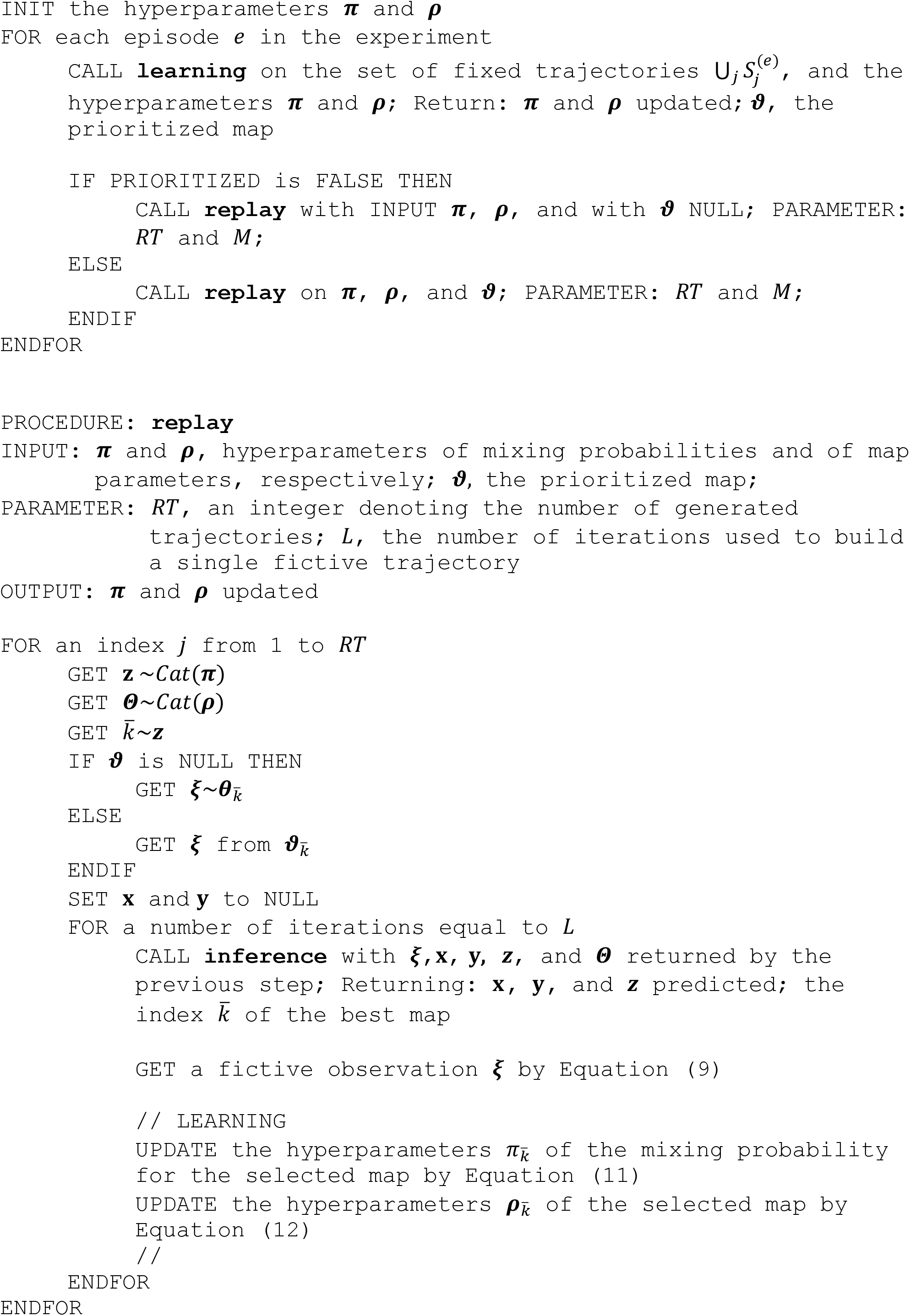

